# A Clinically Useful and Biologically Informative Genomic Classifier for Papillary Thyroid Cancer

**DOI:** 10.1101/2022.12.27.522039

**Authors:** Steven Craig, Cynthia Stretch, Farshad Farshidfar, Dropen Sheka, Nikolay Alabi, Ashar Siddiqui, Karen Kopciuk, Young Joo Park, Moosa Khalil, Faisal Khan, Adrian Harvey, Oliver F. Bathe

**Author notes:** Corresponding Author: Oliver F. Bathe, Division of Surgical Oncology, Tom Baker Cancer Centre, 1331-29^th^ St NW, Calgary, AB, Canada, T2N 4N2.

## Abstract

Clinical management of papillary thyroid cancer depends on estimations of prognosis. Standard care, which relies on prognostication based on clinicopathologic features, is inaccurate. We applied a machine learning algorithm (*HighLifeR*) to 502 cases annotated by The Cancer Genome Atlas Project to derive an accurate molecular prognostic classifier. Unsupervised analysis of the 82 genes that were most closely associated with recurrence after surgery enabled identification of three unique molecular subtypes. One subtype had a high recurrence rate, an immunosuppressed microenvironment, and enrichment of the EZH2-HOTAIR pathway. Two other unique molecular subtypes with a lower rate of recurrence were identified, including one subtype with a paucity of BRAF^V600E^ mutations and a high rate of RAS mutations. The genomic risk classifier, in addition to tumor size and lymph node status, enabled effective prognostication that outperformed the American Thyroid Association clinical risk stratification. The genomic classifier we derived can potentially be applied preoperatively to direct clinical decision-making. Distinct biological features of molecular subtypes also have implications regarding sensitivity to radioactive iodine, EZH2 inhibitors, and immune checkpoint inhibitors.

---

Although there has been a dramatic rise in the incidence of papillary thyroid cancer (PTC) in recent decades ^1–3^, the unique tumor biology of PTC remains poorly understood. PTC is characterized by frequent and early spread to regional lymph nodes, with occasional invasion into surrounding soft tissues. Despite the propensity of PTC to metastasize to locoregional lymph nodes, PTC has a very low incidence of distant metastasis and high overall cure rates. On the other hand, 10 - 15% of PTC cases behave more aggressively and show a greater proclivity for disease recurrence and resistance to conventional adjuvant therapies such as radioactive iodine (RAI)^4^. Understanding what drives this dichotomous clinical behavior is important for two reasons. First, recognizing the biological mechanisms that characterize more aggressive PTC variants may reveal novel therapeutic targets. Second, clinical management can be influenced by accurate prognostication. Patients with a good prognosis can be managed with more conservative surgery or active surveillance, RAI could be avoided, and follow-up regimes could be de-escalated. Conversely, the identification of aggressive PTCs would appropriately direct more extensive surgery, adjuvant therapies, and more intensive or prolonged follow-up periods.

The American Thyroid Association (ATA) Risk Stratification system estimates the risk of disease recurrence based on clinical and pathological factors. It is the most widely used system to estimate prognosis and guide postoperative clinical decision-making^5^. Although the ATA Clinical Risk Stratification system has been validated retrospectively, the proportion of variance explained is suboptimal^6, 7^. The inability of the ATA system to predict recurrence accurately may be because it is insufficiently informed by molecular features. Currently, only one molecular marker, the BRAF^V600E^ gene mutation, is incorporated in the ATA Risk Stratification System^5^, and its value in prognostication is unclear^8^.

In 2014, The Cancer Genome Atlas (TCGA) Network published a landmark study that described the complete genomic landscape of papillary thyroid cancer^9^. A comprehensive description of the molecular features of PTC was provided, and distinct molecular subgroups were identified through the use of unsupervised hierarchical clustering. Two meta-clusters were identified: one consisting of BRAF^V600E^ -driven tumors, and one with RAS-mutated tumors. At the mRNA, miRNA, DNA methylation, and protein expression levels, the number of subtypes varied, but these subtypes were predominantly associated with one of the two meta-clusters. While the study provided insight to the molecular diversity and classification of PTC, it did not relate molecular features with clinical outcomes, such as progression-free survival (PFS).

An alternative approach to deriving molecular classifications of disease is to identify biologically important variants by evaluating their association with clinically relevant phenotypes. For example, PFS is a surrogate marker for the phenotype in that it distinguishes between aggressive and indolent disease. This approach can provide novel insights into the molecular mechanisms that are responsible for the manifestations of the phenotype. *HighLifeR* is a proprietary machine learning algorithm that robustly identifies features of a highly dimensional dataset that are very closely associated with survival outcomes. By applying the *HighLifeR* algorithm to the transcriptional data in the TCGA cohort, we devised a molecular classifier that outperforms the ATA Risk Stratification system in predicting the likelihood of recurrence. Importantly, in PTCs classified by our molecular risk index, we uncovered several molecular features that point to pathogenic mechanisms that could be targeted therapeutically.

## RESULTS

### Identification of robust molecular signatures associated with recurrence risk

*HighLifeR* utilizes a workflow based on the partial Cox proportional hazards regression model to construct independent predictive principal components. The Cox proportional hazards are calculated from an exhaustive series of permutations that have millions of combinations of genes and patients in a structured machine learning environment. We applied the *HighLifeR* algorithm to the RNA-Seq expression dataset from TCGA. This dataset contains batch-corrected expression levels of more than 20,000 genes in a set of 502 PTC patients (335 of whom were used as discovery set). In total, we tested the associations of genes with progression free survival (PFS) in more than 7,500,000 combinations of genes and subsets of the discovery cohort. We identified 44 genes that satisfied our criteria for prognostic significance. Unsupervised clustering based on these 44 genes revealed three molecular subgroups with significantly different PFS (Log rank p = 0.0001) (Figure 1A and 1B). To address the potential effects of censored events, we reapplied the *HighLifeR* algorithm to all cases that had at least 36 months of follow-up with known outcomes (i.e., disease-free after at least 36 months or a recurrence during that time) (*N* =222). An additional set of 41 prognostic genes was identified. Three of these genes overlapped with the first gene set (*EZH2*, *MTMR14*, and *ZNF215*). The complete set of 82 genes (Table S1) was used in classifier development utilizing the three unsupervised clusters (Figure 1A) as the training classifications.

**Figure 1.**
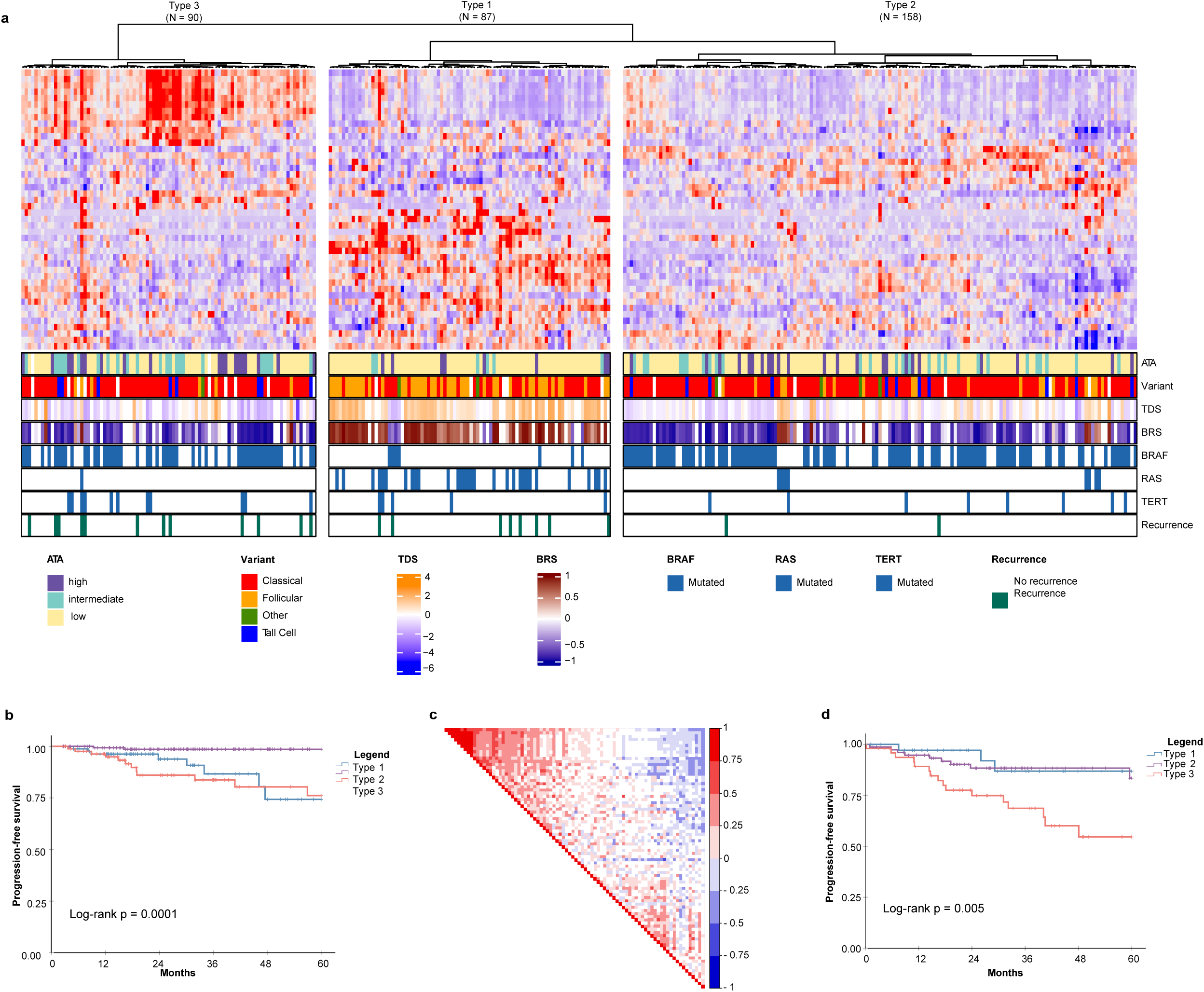
Discovery of mRNA-based prognostic risk groups. **A** Unsupervised clustering of 44 prognostic genes identified by *HighLifeR* shows three distinct clusters in our training set. The heatmap annotation shows distinct differences between molecular subtypes. Specifically, striking differences were in variant type, thyroid differentiation score (TDS), BRAF-RAS score (BRS), and the number of mutations in *BRAF*, *RAS* and *TERT* genes. **B** Survival analysis of these three clusters revealed significant differences in progression-free survival. **C** Correlation matrix associated with all 82 genes identified by HighLifeR shows poor correlation between prognostic genes. **D** Validation of our classification algorithm using TCGA patients excluded from our training dataset (*N* =167) clearly discriminated the three different molecular subtypes and these had significantly different progression-free survival.

Most of the prognostic genes were not correlated with each other. However, there was a cluster of genes with significant positive correlation (Figure 1C). The most highly correlated genes (Pearson r > 0.75, P<0.0001) are involved in mitosis, cell cycle control, and chromatin remodeling. According to The Human Protein Atlas, 49 of the proteins encoded by these 82 genes are detectable by immunohistochemistry in thyroid cancers.

For the purpose of description, the subtypes were labelled as Types 1, 2 and 3. Several interesting features were immediately apparent (Figure 1A). Type 1 PTCs had a paucity of BRAF^V600E^ mutations, were enriched with mutations in NRAS and HRAS, and included most follicular variants. Type 1 PTCs had no cases of tall cell histology; tall cell variants were most common in Type 3. Type 2 and 3 tumors had many of the same clinical features, including BRAF^V600E^ mutations in over half. Type 3 tumors comprised the majority of TERT promoter mutations. Interestingly, the pattern of expression of the top 44 prognostic genes in Type 1 was the inverse of the pattern in Type 3 PTCs (Figure 1A).

A two-step predictive model based on random forest was developed to classify the validation set. The first step identified Type 3 PTCs with a 10-fold cross-validation accuracy of 92%; the second step differentiated Type 1 and 2 PTC with a 10-fold cross-validation accuracy of 86%. As with the discovery cohort, Type 3 PTCs had the highest rate of recurrence (Figure 1D). However, unlike in the discovery cohort, Type 1 tumors had the lowest recurrence rate. We considered that the higher incidence of recurrence could be attributable to disease stage. Indeed, the recurrences in Type 2 PTCs in the validation cohort occurred mostly in patients with advanced tumors (tumors >4 cm *or* N1 disease), which comprised a larger proportion of the validation cohort (61.8%) than the discovery cohort (51.6%).

Univariate and multivariable survival analyses were performed on the discovery set and the validation set. In the larger discovery cohort, factors that were significant on multivariable analysis were molecular subtype, M stage, and presence of tall cell histology (Table 1). In the validation set, molecular subtype and M stage were significant factors in the multivariable analysis (Table S2).

**Table 1.** Univariate and multivariable analysis of factors associated with 5-year PFS in the discovery set.

| Risk Factor | Univariate Analysis |  | Multivariable Analysis |  |
| --- | --- | --- | --- | --- |
|  | Hazard Ratio (95 % CI) | *P* | Hazard Ratio (95 % CI) | *P* |
| Sex (female v. male) | 1.78 (0.75 – 4.26) | 0.19 |  |  |
| Age (< 50 years v. ≥50 years) | 1.86 (0.80 – 4.32) | 0.15 |  |  |
| Tumor size (≤4 cm v. >4 cm) | 1.32 (0.48– 3.61) | 0.59 |  |  |
| Histological type |  |  |  |  |
| Classical | 1 (reference) |  | 1 (reference) |  |
| Follicular | 0.80 (0.23 – 2.77) | 0.72 | 0.27 (0.05 – 1.47) | 0.13 |
| Tall Cell | 4.25 (1.39 – 12.98) | **0.01** | 5.98 (1.29 – 27.72) | **0.02** |
| Other | 0.00 (0.00 – inf) | 1.0 | 0.00 (0.00 – inf) | 1.0 |
| T stage |  |  |  |  |
| T1 | 1 (reference) |  | 1 (reference) |  |
| T2 | 3.62 (0.78 – 16.77) | 0.10 | 4.37 (0.90 – 21.28) | 0.07 |
| T3 | 4.02 (0.85 – 18.92) | 0.08 | 3.10 (0.61 – 15.65) | 0.17 |
| T4 | 7.76 (1.28 – 46.92) | **0.03** | 0.93 (0.08 – 11.34) | 0.95 |
| N stage (N0/NX v. N1) | 1.22 (0.53 – 2.82) | 0.65 |  |  |
| M stage (M0 v. M1) | 8.92 (3.00 – 26.53) | **8.22 x 10<sup>-5</sup>** | 11.02 (1.48 – 82.16) | **0.019** |
| BRAF<sup>V600E</sup> mutation (Absent v. Present) | 1.66 (0.70 – 3.98) | 0.25 |  |  |
| TERT promoter mutation (Absent v. Present) | 5.26 (2.06 – 13.45) | **5.31 x 10<sup>-4</sup>** | 2.13 (0.49 – 9.30) | 0.31 |
| Thyroid Differentiation Score |  |  |  |  |
| < -1 | 1 (reference) |  |  |  |
| -1 to 0 | 0.64 (0.21 – 1.90) | 0.42 |  |  |
| > 0 | 0.46 (0.16 – 1.38) | 0.17 |  |  |
| BRAF<sup>V600E</sup>-RAS class |  |  |  |  |
| BRAF-like | 1 (reference) |  |  |  |
| RAS-like | 1.15 (0.44 – 3.01) | 0.77 |  |  |
| ATA Risk |  |  |  |  |
| Low | 1 (reference) |  | 1 (reference) |  |
| Intermediate | 3.13 (1.11 – 8.81) | **0.03** | 1.71 (0.37 – 7.96) | 0.49 |
| High | 3.34 (1.24 – 8.99) | **0.02** | 1.90 (0.42 – 8.61) | 0.40 |
| Molecular Subtype |  |  |  |  |
| Type 2 | 1 (reference) |  | 1 (reference) |  |
| Type 1 | 9.40 (1.99 – 44.46) | **0.005** | 17.98 (3.27 – 98.92) | **8.97 x 10<sup>-4</sup>** |
| Type 3 | 12.93 (2.89 – 57.80) | **8.1 x 10<sup>-4</sup>** | 9.77 (2.04 – 46.81) | **0.004** |

HR = Hazard ratio. This is the ratio of recurrence rates in two comparator groups for any independent variable (for binary variables) or the logarithm of the change in death rate per unit change of the independent variable (if the variable is continuous).
CI = Confidence interval

We considered that anaplastic thyroid cancer (ATC), an extremely aggressive form of thyroid cancer with a high incidence of TERT promoter mutations, may arise from well differentiated thyroid cancer ^10^. Our molecular classification algorithm was applied to RNASeq data from 10 cases of ATC arising from PTC (Bioproject ID: PRJNA 523137; https://www.ebi.ac.uk/ena/browser/view/PRJNA523137)^11^. Interestingly, all 10 cases were clearly classified as the Type 3 molecular subtype.

### Clinicopathological features of molecular subtypes

To comprehensively describe the clinical and molecular differences between the three molecular subtypes, we combined the discovery and validation cohorts. Clinical features are summarized in Table 2. Altogether, 25.0% were Type 1, 47.3% were Type 2, 27.7% were Type 3 PTCs. There was no significant relationship between molecular subtype and age, sex, race, or ethnicity. The incidence of multifocality was the same in each molecular subtype. Extrathyroidal spread and lymph node metastases were significantly more prevalent in Type 2 and 3 PTCs. Type 1 tumors tended to have a lower T-stage and had the lowest incidence of lymph node metastases.

**Table 2.** Clinical characteristics for the three molecular subtypes - for training and test set samples combined.

| Factor | Molecular subtype |  |  | P |
| --- | --- | --- | --- | --- |
|  | Type 1 (N = 126) | Type 2 (N = 237) | Type 3 (N = 139) |  |
| **Sex, N (%)** |  |  |  |  |
| Male | 35 (27.8) | 66 (27.8) | 35 (25.2) | 0.84<sup>a</sup> |
| Female | 91 (72.2) | 171 (72.2) | 104 (74.8) |  |
| **Age, mean ± SD** | 49.5 ± 15.3 | 46.8 ± 14.3 | 46.1 ± 18.4 | 0.14<sup>b</sup> |
| **Race, N (%)** |  |  |  | 0.74<sup>a</sup> |
| White | 69 (79.3) | 160 (81.2) | 102 (81.0) |  |
| Black or African American | 8 (9.2) | 12 (6.1) | 7 (5.6) |  |
| American Indian or Alaskan Native | 0 (0.0) | 0 (0.0) | 1 (0.8) |  |
| Asian | 10 (11.5) | 25 (12.7) | 16 (12.7) |  |
| **Ethnicity, N (%)** |  |  |  | 0.58<sup>a</sup> |
| Hispanic or Latino | 6 (7.0) | 17 (9.1) | 14 (11.2) |  |
| Not Hispanic or Latino | 80 (93.0) | 170 (90.9) | 111 (88.8) |  |
| **Histological type, N (%)** |  |  |  |  |
| Classical | 55 (43.7) | 187 (78.9) | 112 (80.6) | < 0.0001<sup>*a</sup> |
| Follicular | 67 (53.2) | 24 (10.1) | 10 (7.2) |  |
| Tall Cell | 0 (0.0) | 21 (8.9) | 15 (10.8) |  |
| Other | 4 (3.2) | 5 (2.1) | 2 (1.4) |  |
| **Focality, N (%)** |  |  |  | 0.38<sup>a</sup> |
| Unifocal | 67 (54.0) | 119 (51.3) | 80 (58.8) |  |
| Multifocal | 57 (46.0) | 113 (48.7) | 56 (41.2) |  |
| **Extrathyroidal spread, N (%)** |  |  |  | < 0.0001<sup>*a</sup> |
| None | 105 (88.2) | 141 (61.0) | 86 (64.2) |  |
| Minimal | 13 (10.9) | 82 (35.5) | 39 (29.1) |  |
| Gross (T4a and T4b) | 1 (0.8) | 8 (3.5) | 9 (6.7) |  |
| **T Stage, N (%)** |  |  |  |  |
| T1 | 40 (31.7) | 65 (27.4) | 39 (28.1) | 0.003<sup>*a</sup> |
| T2 | 56 (44.4) | 68 (28.7) | 41 (29.5) |  |
| T3 | 28 (22.2) | 94 (39.7) | 49 (35.3) |  |
| T4 | 2 (1.6) | 10 (4.2) | 10 (7.2) |  |
| **N Stage, N (%)** |  |  |  | < 0.0001<sup>*a</sup> |
| N0/NX | 104 (82.5) | 118 (49.8) | 58 (41.7) |  |
| N1 | 22 (17.5) | 119 (50.2) | 81 (58.3) |  |
| **M Stage, N (%)** |  |  |  | 0.93<sup>a</sup> |
| M0 | 123 (98.4) | 233 (98.3) | 136 (97.8) |  |
| M1 | 2 (1.6) | 4 (1.7) | 3 (2.2) |  |
| **RAS mutation, N (%)** |  |  |  | < 0.0001<sup>*a</sup> |
| Absent | 82 (67.2) | 225 (95.7) | 132 (98.5) |  |
| Present | 40 (32.8) | 10 (4.3) | 2 (1.5) |  |
| **BRAF<sup>V600E</sup> mutation, N (%)** |  |  |  |  |
| Absent | 118 (94.4) | 86 (36.3) | 64 (46.0) | < 0.0001<sup>*a</sup> |
| Present | 7 (5.6) | 151 (63.7) | 75 (54.0) |  |
| **TERT promoter mutation, N (%)** |  |  |  |  |
| Absent | 117 (94.4) | 225 (95.7) | 118 (86.1) | 0.002<sup>a</sup> |
| Present | 7 (5.6) | 10 (4.3) | 19 (13.9) |  |
| **Thyroid Differentiation Score, N (%)** |  |  |  | < 0.0001<sup>a</sup> |
| < -1 | 3 (3.0) | 39 (21.3) | 38 (37.3) |  |
| -1 to 0 | 6 (6.1) | 81 (44.3) | 37 (36.3) |  |
| > 0 | 90 (90.9) | 63 (34.4) | 27 (26.5) |  |
| **BRAF<sup>V600E</sup>-RAS score, mean ± SD** | 0.67 ± 0.43 | -0.65 ± 0.45 | -0.60 ± 0.49 | < 0.0001<sup>b</sup> |
| **ATA Risk, N (%)** |  |  |  | < 0.0001<sup>a</sup> |
| Low | 106 (84.8) | 146 (61.9) | 74 (53.2) |  |
| Intermediate | 7 (5.6) | 51 (21.6) | 40 (28.8) |  |
| High | 12 (9.6) | 39 (16.5) | 25 (18.0) |  |
| **AMES, N (%)** |  |  |  | 0.06<sup>a</sup> |
| Low | 113 (90.4) | 198 (83.9) | 111 (79.9) |  |
| High | 12 (9.6) | 38 (16.1) | 28 (20.1) |  |
| **MACIS score, N (%)** |  |  |  | 0.054<sup>a</sup> |
| < 6.00 | 84 (71.2) | 182 (81.6) | 99 (76.2) |  |
| 6.00 to 6.99 | 20 (16.9) | 22 (9.9) | 13 (10.0) |  |
| 7.00 to 7.99 | 12 (10.2) | 12 (5.4) | 9 (6.9) |  |
| > 8.00 | 2 (1.7) | 7 (3.1) | 9 (6.9) |  |

a = Pearson's Chi-squared test
b = Kruskal–Wallis test
\* = significant

We explored the relationships of prognostic groups and molecular subtypes with ATA risk, AMES score and MACIS score. ATA risk classes were more frequently low risk in Type 1 PTCs, and high-risk ATA class was more frequent in Type 2 and 3 PTCs. A high AMES score (corresponding to a higher risk of death from PTC) was least frequent in Type 1 tumors and most frequent in Type 3 tumors. Similarly, high MACIS scores were most frequent in Type 3 PTCs. In all, clinicopathological features in Type 2 and Type 3 PTCs were not easily distinguishable, but Type 1 PTCs were markedly different.

Two gene signatures described by TCGA^9^, the BRAF^V600E^-RAS score (BRS) and the Thyroid Differentiation Score (TDS), were evaluated in the context of the molecular subtypes. Importantly, only two of the genes from our signature overlapped with the genes that comprise BRS: CTSC and FN1; and none of the 82 genes overlapped with genes that comprise TDS. The BRAF^V600E^-RAS score (BRS), which quantifies the similarity of the gene expression profile to either BRAF^V600E^ or RAS mutant profiles (−1 to +1) ^9^, was significantly higher in Type 1 PTCs (Table 2). Using this method of classification, 91.9% of Type 1 PTCs were RAS-like with a corresponding higher incidence of NRAS and HRAS mutations. In contrast, 91.8% of Type 2 PTCs and 88.2% of Type 3 PTCs were BRAF-like. The Thyroid Differentiation Score (TDS, range = −4.08 to 2.59) decreased with a higher histological grade^9^ and was lowest in the Type 3 molecular subgroup (Table 2, Figure 1A). Neither BRS nor TDS had a significant association with PFS.

Finally, molecular subtypes had distinct mutation patterns. Mutations associated with each molecular subtype are summarized in Figure S1 A – E. NRAS mutations were present in 33.1% of Type 1 PTCs, in 4.3% of Type 2 PTC’s and 1.5% of Type 3 PTCs. There was also a high prevalence of mutations in the thyroglobulin gene (TG) in Type 1 PTCs. BRAF^V600E^ mutations were found in only 5.6% of Type 1 PTCs, but in more than half of Type 2 and 3 PTCs. Although TERT promoter mutations were present in all molecular subtypes, the majority were found in Type 3 PTCs. When we compared Type 2 versus Type 3 tumors, we found no significant differences in mutations.

### Biological features of Type 3 PTCs

To delineate the biological features of each tumor type, molecular features were identified that distinguished Type 3 PTCs from the other two subtypes. In comparison to Type 1 PTCs, there were 2234 differentially expressed genes (DEGs). We performed Gene Set Enrichment Analysis (GSEA) of the 50 hallmark gene sets in MSigDB ^12^. Pathways that were positively enriched were involved in inflammation and epithelial-mesenchymal transition (EMT) (Figure 2). Compared to Type 2 PTCs, there were relatively fewer DEGs (496 DEGs). GSEA demonstrated mostly enrichment in inflammatory pathways, pathways involved in proliferation, as well as mTORC1 signaling (Figure 2). Mindful that the Forkhead box M1 (FOXM1) transcription factor encourages migration and invasion of PTC cells ^13^, we performed a targeted GSEA of the FOXM1 pathway. This pathway was positively enriched in Type 3 cancers in comparison to Type 1 (False Discovery Rate (FDR) =0.002) and Type 2 PTCs (FDR=0.004), as previously reported ^14^.

**Figure 2.**
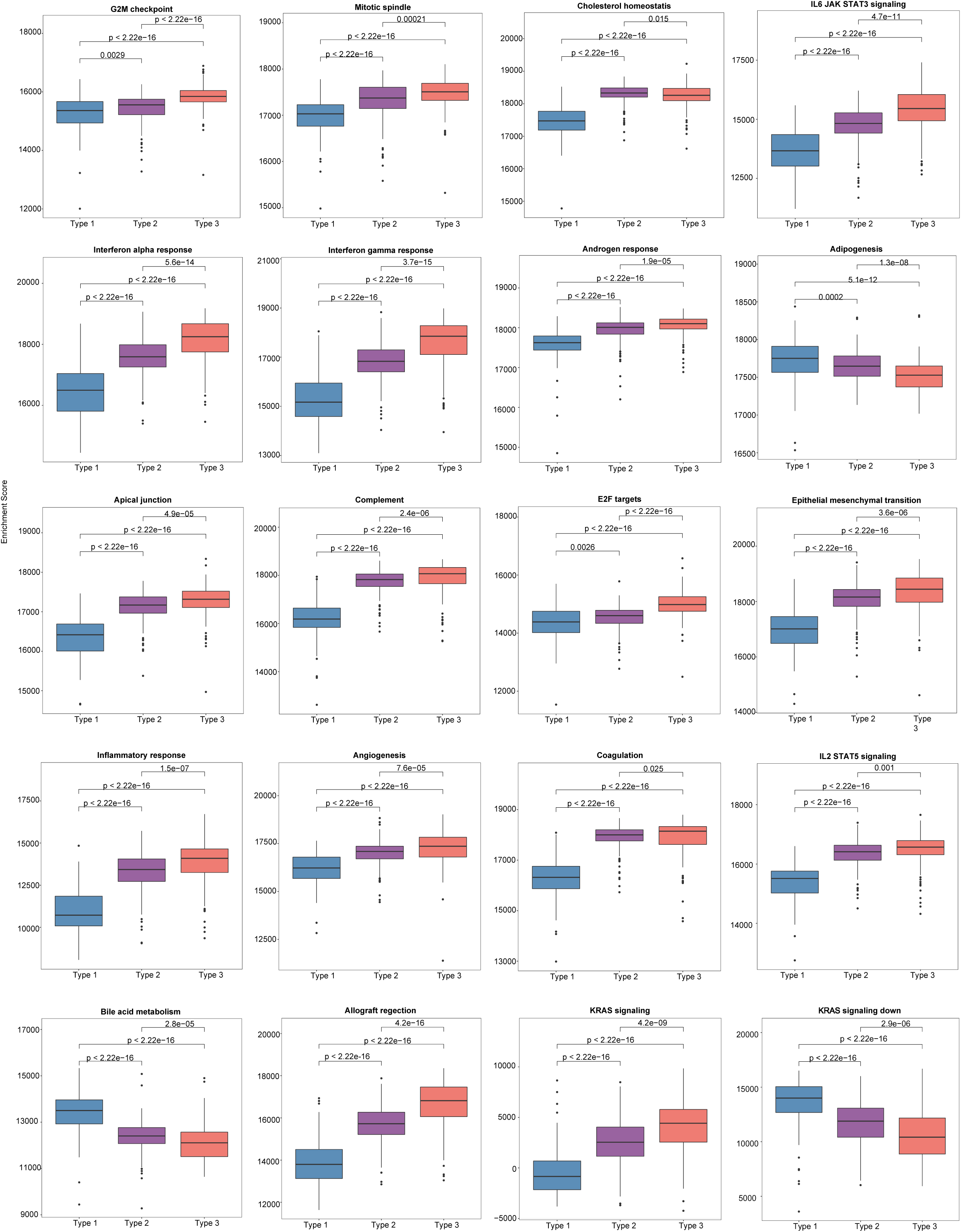
The single-sample gene set enrichment analysis (ssGSEA) scores between different molecular subtypes using the Hallmarks geneset collection. Only the top 20 significantly enriched genesets are shown.

Analysis with Ingenuity Pathway Analysis (IPA) demonstrated intriguing inflammatory features in Type 3 tumors. In comparison to Type 1 PTCs, there was significant enrichment of genes involved in HMGB1 signaling, STAT3 signaling, IL-23 signaling, and IL-17 signaling (Figure S2A). HMGB1 upregulation and the successive overexpression of IL-23, IL-17 and IL-6, followed by STAT3 activation, promotes tumor growth ^15^. STAT3 from tumor cells and myeloid cells is also known to induce IL-23 production by tumor associated macrophages; regulatory T cells expressing IL-23R are then activated to create an immunosuppressive tumor microenvironment^16^. In comparison to Type 2 PTCs, Type 3 showed enrichment of genes involved in HMGB1 signaling and IL-17 signaling, as well as immunosuppressive processes.

Deconvolution of immune cell types using CIBERSORT revealed that Type 3 tumors contained significantly more activated CD4+ T cells, and a high number of CD4+CD25+ regulatory T cells (Figure 3A). The expression levels of immunoregulatory genes were markedly elevated in comparison to the other two molecular subtypes (Figure 3B).

**Figure 3.**
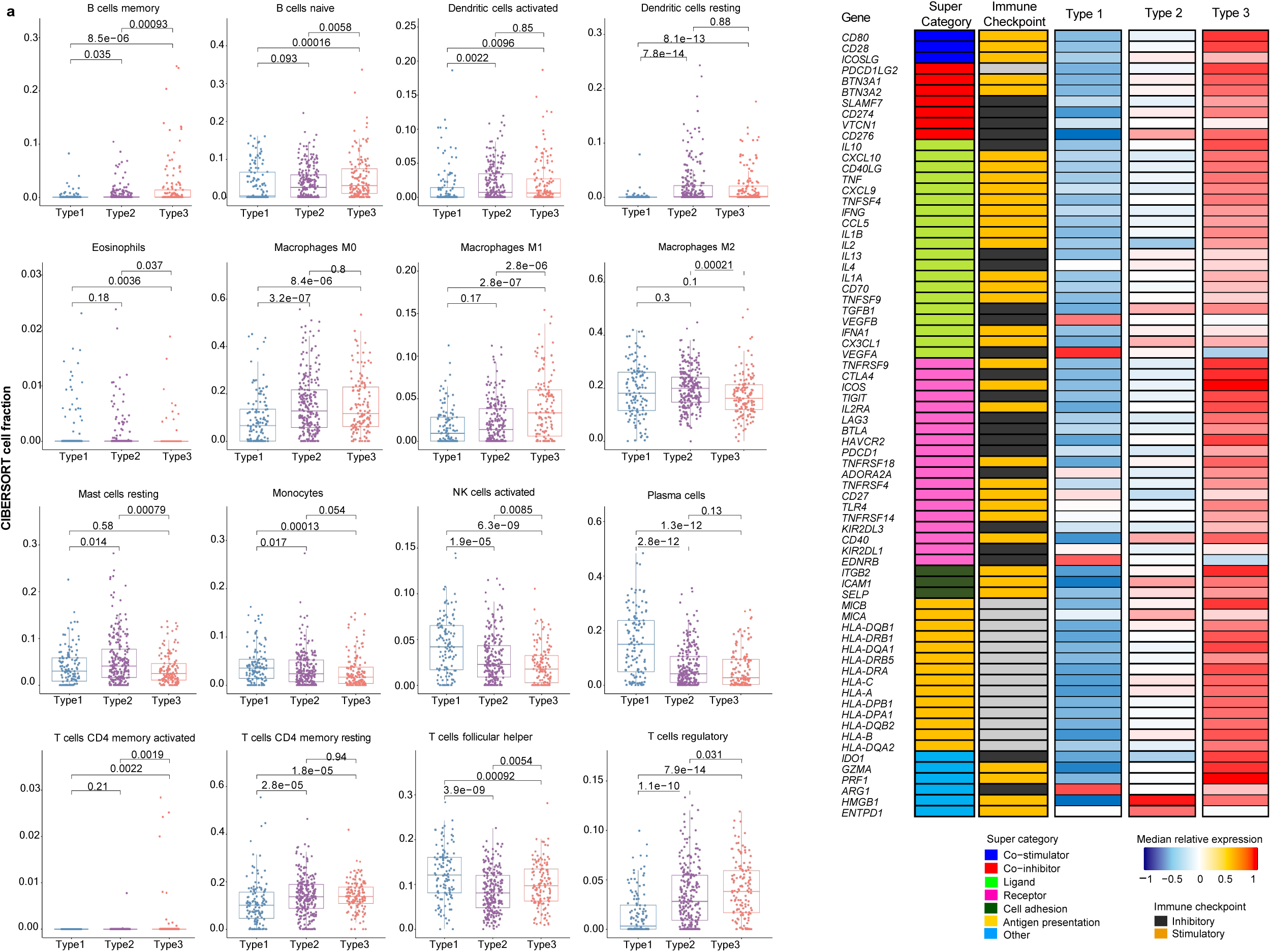
Immune profile of mRNA-based prognostic risk groups. **A** Estimates of immune cell type fractions present in the mRNA-based prognostic risk groups based on deconvolution of expression profiles using CIBERSORT. Immune cell fractions were compared using one-way ANOVA test and p-values are shown for each comparison. Displayed are the 16 of 22 cell types which showed significant differences based on one-way ANOVA. **B** Gene expression of immunomodulators was examined across mRNA-based prognostic risk groups. The immunomodulators were classed into one of seven categories (co-stimulator, co-inhibitor, ligand, receptor, cell adhesion, antigen presentation or other) and each of the immunomodulators were also categorized as immune checkpoint inhibitors or stimulators. Median normalized expression levels for each mRNA-based prognostic risk group are shown.

Another striking feature uncovered in the IPA analysis was the positive enrichment of the HOTAIR (HOXA transcript antisense RNA) pathway. HOTAIR is a lncRNA that interacts with Polycomb Repressive Complex 2 (PRC2), a histone methyltransferase that affects epigenetic silencing in diverse proneoplastic processes including EMT^17^. HOTAIR interaction with PRC2 drives EZH2-mediated gene repression. Elevated EZH2 expression is characteristic of the high-risk phenotype, as is upregulation of HOTAIR (*P* < 0.001, FDR< 0.001). HOTAIRM1, which similarly interacts with EZH2 and encourages an immunosuppressive microenvironment ^18, 19^, was also upregulated (FDR=0.001).

In comparison to Type 1 PTCs, Type 3 PTCs contained 596 differentially methylated genes: 35 were hypermethylated in association with downregulation, and 236 were hypomethylated with coincidental upregulation at the mRNA level (Figure S3A). There were no differentially methylated genes between Type 3 and Type 2 PTCs. There were 571 differentially expressed (DE) miRNAs in comparison to Type 1 PTCs. There were 85 DE miRNAs compared to Type 2 PTCs. In sum, differences between Type 3 and Type 2 were not pronounced. The epigenetic features and the miRNA expression pattern appeared to drive the inflammatory and immunosuppressive features of Type 3 PTCs (Figure S3C and Figure S4A).

### Biological Features of Type 2 PTCs

In comparison with Type 1 PTCs, there were 1124 DEGs in Type 2 PTCs. As in Type 3 tumors, GSEA of the 50 hallmark pathways demonstrated significant enrichment in proinflammatory gene sets and genes involved in EMT (Figure 2). A targeted GSEA of the FOXM1 pathway demonstrated positive enrichment (*P* = 0.007), although this was less pronounced than in Type 3 PTCs. IPA demonstrated enrichment in EMT factors, as well as IL-6 and IL-17 signaling (Figure S2B). Immunosuppressive processes were notably absent, consistent with the lower expression levels of immunoregulatory genes in Type 2 PTCs compared to Type 3. The HOTAIR pathway was positively enriched but was less pronounced than in Type 3 PTCs.

In comparison to Type 1 PTCs, Type 2 PTCs contained 600 differentially methylated genes: 10 were hypermethylated in association with downregulation, and 240 were hypomethylated with coincidental upregulation at the mRNA level (Figure S3B). While epigenetic changes could be linked to some nonspecific inflammatory changes (Figure S3D), functional differences between Type 2 and Type 1 PTCs appeared to be more attributable to differences in miRNA expression. Relative to Type 1 PTCs, there were 169 upregulated miRNAs with downregulated mRNA targets, and 218 downregulated miRNAs with upregulated mRNAs targets; and genes so affected were involved in inflammation, the HOTAIR regulatory pathway, and neuroendocrine functions (Figure S4B).

### Biological Features of Type 1 PTCs

Type 1 PTCs are characterized by a very different inflammatory microenvironment compared to Type 2 and 3 PTCs; for example, the fraction of monocytes and activated NK cells in Type 1 PTCs was 148% and 201% greater than Type 3 PTCs, respectively (Figure 3A). On GSEA, there is enrichment of fatty acid metabolism. Bile acid metabolism is also prominent, although it did not reach significance following correction for multiple comparisons (Figure 2). Compared to Type 2 and 3 tumors, IPA based on DEGs demonstrated LXR/RXR activation (Figures S2A and S2B). PTEN signaling and PPAR signaling was also evident in comparison to Type 3 PTCs. These mostly metabolic features appear to be driven mostly by differences at the miRNA level (Figures S4A and S4B). Others have previously observed alterations in lipid metabolism in PTC, although it is unclear how diverse this characteristic is ^20, 21^.

### External validation of molecular subtypes

RNA-Seq expression data from an ethnically and racially distinctive cohort from Seoul National University College of Medicine, Korea ^22^ were evaluated to understand the generalizability of the subgroups we identified. The cohort comprised 48 follicular variant and 76 classical PTCs (total *N* =124). No tall cell variant tumors were included. Similar patterns of expression in the 82 prognostic genes were identifiable. Some of the features that characterized each molecular subtype were seen in the Korean cohort (Table S3). Specifically, Type 1 PTCs had the highest frequency of follicular variants, were mostly RAS-like, and had the highest incidence of RAS mutations. Type 1 PTCs had the lowest incidence of BRAF^V600E^ mutations, although the frequency was higher than in the TCGA cohort. Type 2 and 3 PTCs had the highest incidence of extrathyroidal spread and tended to have a higher incidence of lymph node metastases. Incidence of lymph node disease was lowest in Type 1 PTCs, although this was not significant. The median follow-up was 88 months, and there was only one structural recurrence in an advanced Type 2 PTC. There were 9 biochemical recurrences defined by TSH-stimulated thyroglobulin ≥1 µg/L. Biochemical recurrences occurred in one Type 1 PTC (2.4%), four Type 2 PTCs (5.8%), and four Type 3 PTCs (28.6%). Dates to recurrence events were unknown for this dataset.

To further explore the clinical utility of the molecular risk stratification, we designed an assay based on targeted hybrid-capture enrichment RNA sequencing (RNASeq). Expression of the 82 genes that distinguished molecular classes were quantified in frozen samples from Edmonton, Canada (*N* =132). The median follow-up period for these cases was 66 months. Cases were classified in a way that blinded them to clinical features and outcomes. 5-year recurrence rates for the study cohort are summarized in Table S4. Strikingly, recurrence rates were consistently highest in Type 3 PTCs, regardless of whether tumors were early (tumor size 1-4 cm *and* N0/NX) or advanced. In contrast to Type 3 PTCs, and in keeping with what was found in the TCGA dataset, Type 1 and Type 2 PTCs recurrence rates were higher in advanced tumors.

### Potential Clinical Utility of Molecular Risk Stratification

While Type 3 PTCs consistently had a worse prognosis, given the instability of recurrence outcomes of Type 1 and Type 2 PTCs in the TCGA discovery and validation cohorts, we considered that clinical factors could modify risk. Two factors that can be readily evaluated prior to surgery are tumor size and lymph node involvement. Indeed, according to the 2015 ATA guidelines, patients with the following *preoperative* criteria are considered candidates for thyroid lobectomy: tumor size 1 - 4 cm, clinically node negative, no extrathyroidal spread, no family history of thyroid cancer, and no history of radiation therapy to the neck ^5^. Using the combined data from the TCGA cohort, the Korean cohort and the Edmonton cohort (*N* =743), it was apparent that the lowest risk for recurrence was in Type 1 and Type 2 PTCs with early PTCs (tumor size ≤4 cm and no lymph node disease). A higher risk was observed in advanced Type 1 and 2 PTCs (tumor size >4 cm or N1) The highest risk of recurrence was in Type 3 PTCs, regardless of whether early or advanced (Figure 4A). Altogether, this meant by considering both the molecular subtype and the clinical information (tumor size >4 cm or N1) we could describe two distinct risk classes (a high- and a low-risk class) (Figure 4B).

**Figure 4.**
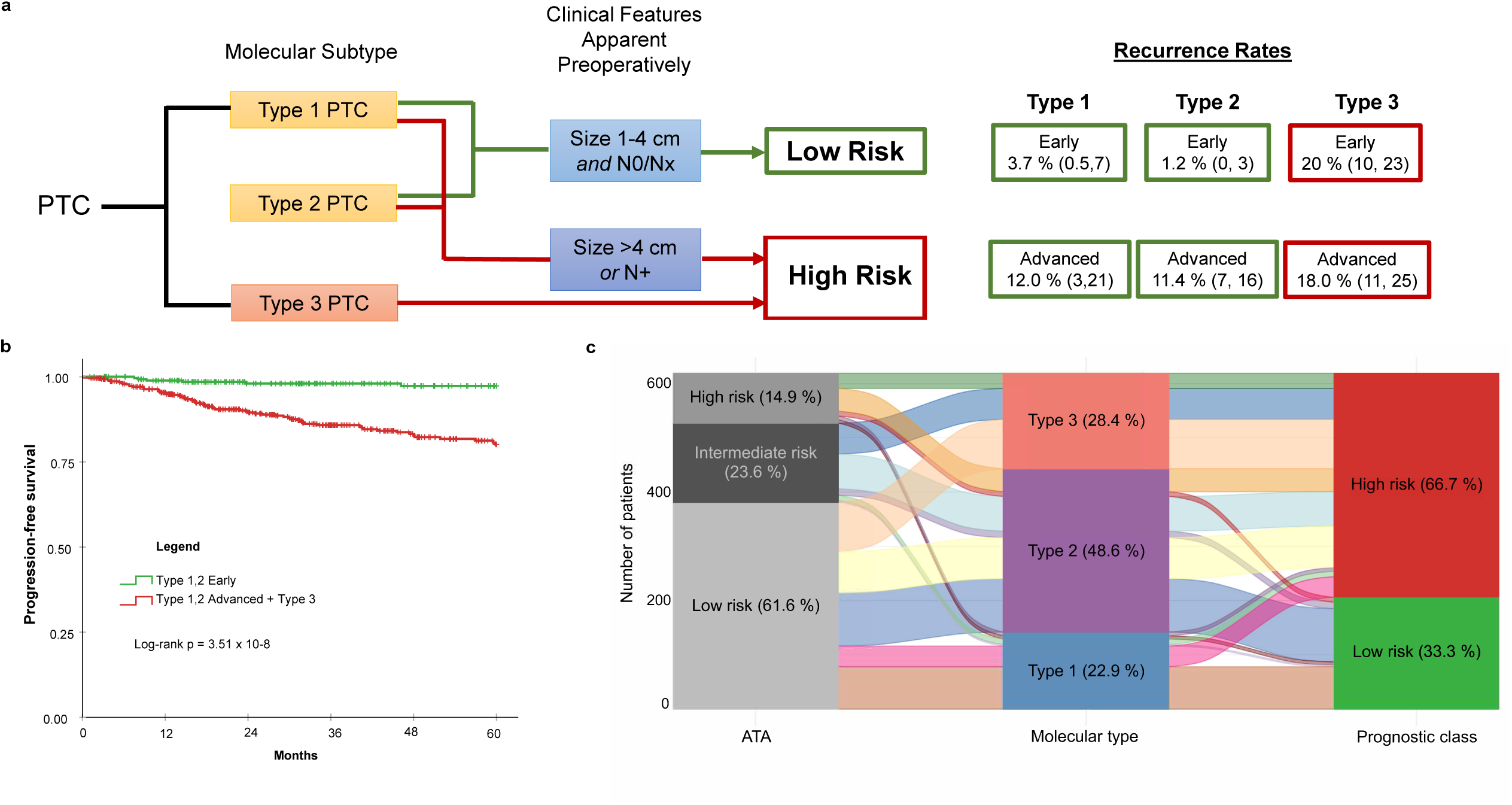
Molecular subtype in conjunction with two simple clinical factors improves recurrence risk stratification for PTC. **A** Proposed risk stratification flow diagram which incorporates molecular subtype determination, followed by assessment of tumor size and lymph node status to ultimately determine if tumors have a low or a high risk of recurrence. **B** Dichotomization resulting from incorporation of molecular subtype with preoperatively apparent clinical features shows significantly different survival (Log rank *P* = 3.51 x 10-8). The low-risk class exhibits very few recurrences. **C** Alluvial plot illustrates how patients originally assigned into the low-, intermediate-, and high-risk groups using ATA are re-classified into the three molecular subtypes and subsequently into the two risk classes after incorporating clinical features (i.e., tumor size and lymph node status).

Molecular subtyping in conjunction with tumor size and lymph node status can simplify risk stratification in comparison to the ATA Risk Stratification System (2015) ^5^. The ATA Risk Stratification System is the most commonly used clinical risk index for predicting disease recurrence for differentiated thyroid cancer. Performance of the two risk stratification methods was compared. A consensus ATA risk classification for each case was established by two practicing clinicians. To calculate 5-year time-dependent AUROC, a binary classification was applied: low risk vs. intermediate/high risk. Data from the Korean cohort were not included because of the paucity of recurrence events. Using the smoothROCtime package^23^, AUROC was 0.80 for molecular classification + tumor size/lymph node status; 0.70 for molecular subtype alone; and 0.51 for the ATA risk stratification method. This is not surprising considering the low rate of recurrence for the molecular low-risk prognostic class (i.e., Type 1 or Type 2, size 1-4 cm and N0/Nx PTCs) and the redistribution of class assignments between the ATA risk classes and prognostic class (Figure 4C): Twenty-three of the 205 ATA low risk PTCs were reclassified to high risk, and 21 of those had recurrences.

The most impactful application of our findings is likely to be in the preoperative period. One problem with using purely clinical criteria to select lobectomy candidates is that 40 - 60% will require a completion thyroidectomy because the *postoperative* risk stratification classified them as a higher risk than initially estimated^24–29^. If molecular subtype could be reliably determined by fine needle aspirate (FNA), patient selection for more conservative surgery could be refined. In the same way, for tumors measuring <1 cm, patients could be selected for active surveillance with greater assurance.

### Potential therapeutic approaches for aggressive PTCs

In addition to the potential application of the prognostic biomarker for surgical decision making, the novel molecular subtypes identify phenotypes that could guide future therapeutic approaches and clinical trials. Radioactive iodine (RAI) following a total thyroidectomy is a conventional approach to ATA classified intermediate- and high-risk tumors. This approach could be modified based on molecular subtypes. For example, TDS is derived from expression levels of 16 thyroid function genes. One such gene, SLC55A5, encodes sodium iodide symporter and is required for iodine uptake by thyrocytes. A high TDS would therefore be most susceptible to RAI. Others have reported that BRAF^V600E^ mutations decrease expression of sodium iodide symporter, thought to be a mechanism of radioactive iodine resistance ^30, 31^. Indeed, in this series, cases with BRAF^V600E^ mutations were associated with lower TDS, regardless of molecular subtype. However, TDS appeared more related with molecular subtype than BRAF^V600E^ mutation status – Type 1 tumors had significantly higher TDS than Type 2 or Type 3 tumors (Figure S5A). Moreover, low TDS was associated with a shorter PFS in patients who received RAI (p < 0.001), but not in patients who did not receive RAI. In the case of ATA intermediate-risk tumors, there have been disparate results in clinical series reporting the benefits of RAI in intermediate-risk PTC ^32, 33^. This can be explained by the differences in TDS between (and within) molecular subgroups (Figure S5A).

There has been substantial interest in pharmacological approaches to restoring tumor differentiation and sensitivity to RAI. MAPK inhibitors have been found to restore expression of thyroid-specific genes and sensitivity to RAI ^31 34^. A phase 2 clinical trial of selumetinib reversed radioiodine resistance in patients with advanced thyroid cancer with clinical benefit ^35^. Rather than focusing on tumors with the BRAF^V600E^ mutation, trials could focus on tumors with a low TDS and a particular molecular subtype. For example, Type 3 PTC overexpress EZH2, a master regulator of chromatin (Figure 5A). EZH2 hypertrimethylation of histone H3 lysine 27 (H3K27) leads to cancer cell de-differentiation. Indeed, a lower TDS is associated with a higher EZH2 expression (Figure 5B). Preclinical studies have demonstrated that EZH2 inhibitor tazemetostat in combination with MAPK inhibitors promotes ^125^Iodine uptake and enhanced cytotoxicity in PTC cells ^34^.

**Figure 5.**
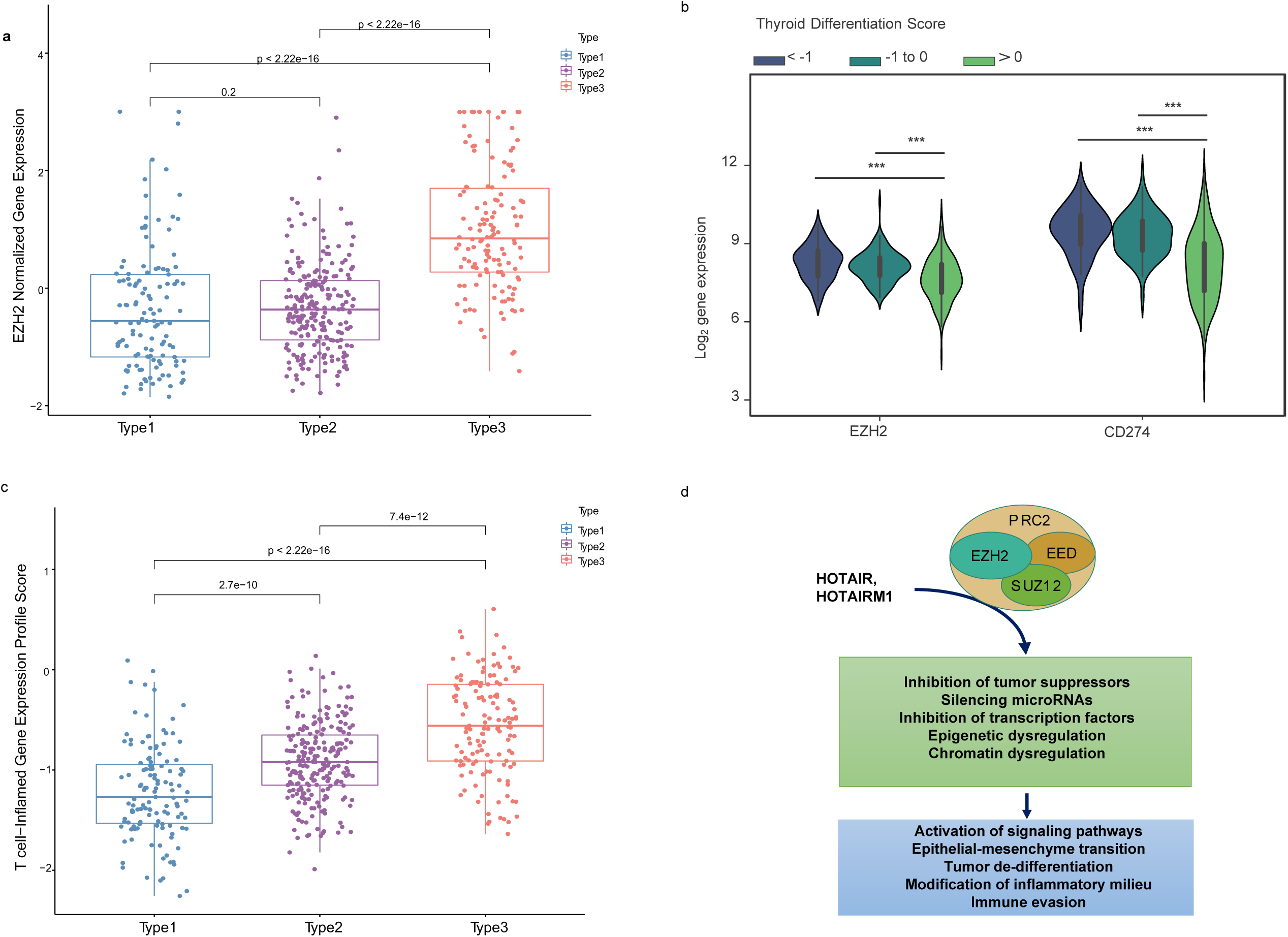
Biological differences between mRNA-based prognostic risk indicates potential therapeutic approaches for aggressive papillary thyroid carcinomas. **A** The relationship between molecular subtype and EZH2 expression shows a higher expression of EZH3 in Type 3 tumors. **B** The relationship between thyroid differentiation score (TDS) and expression of *EZH2* and *CD274* (gene name for PD-L1) are shown in the violin plot. One-way ANOVA p-value <0.0001 is indicated by ***. **B** T-cell-inflamed gene expression scores for each molecular subtype. T-test p-values are shown for each comparison. **C** Diagram of suggested role of EZH2 and associated molecules contributing to the high-risk group phenotype suggests EZH2 inhibitors may favorably modify high-risk tumors.

Type 3 PTCs, characterized by an abundant immune cell infiltrate and higher expression of immunoregulatory molecules (Figure 3A, B), may also be amenable to immune checkpoint inhibitors. In 2018, it was reported that higher T cell-inflamed gene expression profile (GEP) scores were a prerequisite for clinical benefit from PD-1 blockade ^36^. Type 3 tumors identified by our biomarker had the highest T cell-inflamed GEP scores (Figure 5C). In the phase 1b KEYNOTE-028 trial, 22 patients with advanced papillary follicular thyroid cancers received pembrolizumab, and anti-PD-1 antibody ^37^. The overall response rate was only 9.1% and the stable disease rate was 54.5%. The classification we described could improve selection of candidates for this approach. EZH2 inhibitors can also favorably modify the immune microenvironment. PTCs with a low TDS are associated with higher expression of EZH2 as well as CD274, which encodes PD-L1 (Figure 5B). As with immunotherapy in general, Type 3 PTCs have the characteristics for this therapy (Figure 5D).

## DISCUSSION

In 2014 the TCGA reported a seminal description of the molecular features of PTC, providing numerous observations never before reported ^9^. Molecular subclassification at various biological levels was accomplished using unsupervised methods. Combining these features (i.e., mutation, methylation, mRNA) enabled identification of meta-clusters. However, with the exception of finding subgroups that displayed the signaling consequences of BRAF^V600E^ mutations and RAS mutations, limited functional information could be derived using this approach. An alternative approach is to link molecular features with phenotypical features including clinical aggressivity.

Two obstacles exist in studying PTC. First, PTC has a good prognosis and only a small proportion of tumors recur after resection. Second, the methods for linking highly dimensional features to continuous variables (such as survival) have not been well described. Using a machine learning algorithm designed to circumvent these problems, genes most consistently associated with recurrence were identified. Submitting those genes to conventional unsupervised methods of classification facilitated identification of unique molecular subtypes.

Prognostic biomarker development has primarily relied on Cox Proportional Hazard (PH) analysis. However, applying Cox PH to a highly dimensional molecular profiling dataset (e.g., a dataset consisting of over 20,000 mRNA transcripts) would result in a highly overfitted model. Another problem with using a Cox PH model on genomic data is the key assumption that proportional hazard functions remain proportional over time. Assigning a linear risk score function to genes may not be valid, as fluctuations in the survival risk in the range of expression can occur^38^. Another issue with biomarker development is that the random splitting of a single cohort into discovery and validation (test) cohorts can potentially result in the spurious identification of genes that are a product of the composition of the discovery cohort. *HighLifeR* addresses these challenges by testing the effects of many permutations of genes in numerous virtual cohorts, ranking genes by their strength of association with the survival outcome and by the consistency of their effect. As a result, it is well suited for identifying molecular features that have a significant relationship with survival outcomes in highly dimensional genomic datasets. Another machine learning algorithm (EACCD) has previously been employed to identify prognostic groups in thyroid cancer using clinical variables^39^. The EACCD algorithm relies on categorical variables as input. When continuous variables are encountered (e.g., tumour size), arbitrary cut-offs must be utilized, limiting the algorithm’s usefulness for identifying prognostic genes.

Several other groups have interrogated the TCGA cohort, utilizing series of Cox Proportional Hazards regression survival analyses in an attempt to identify prognostic gene signatures for PTC. One study identified 38 genes significantly associated with PTC progression, and 24 of these genes were related to the FOXM1 signaling pathway ^14^. Six of the genes identified by that group overlapped with our gene set (TTK, EZH2, SKA3, KIF4A, HIST2H2BF, BUB1). We also observed positive enrichment of the FOXM1 pathway in Type 3 and Type 2 PTCs. The 5-year time-related AUROC was 0.72. Two groups describing panels of 5 prognostic genes (none of which are present in our gene list) had AUROCs of 0.59 and 0.75 ^40, 41^. Finally, Yang et al. reported on a risk score based on immune infiltrate determined by the Cibersort algorithm^42^. Like us, they found an association of immune checkpoint gene expression and poor prognosis. The AUROC using their approach was 0.71. The strength of our approach is the relatively favourable degree of prognostic accuracy (AUROC 0.80). Moreover, the subtypes we have identified provide meaningful and actionable biological insight by describing potential therapeutic targets.

Importantly, we have described how our prognostic signature can be leveraged with two simple clinical data points: tumor size and lymph node status (used to dichotomize early and advanced PTC). The gene signature by itself performs favorably in comparison to other prognostic gene signatures ^42^. In conjunction with the clinical dichotomization, a subgroup with a low risk of recurrence is identified with greater specificity than the current ATA risk stratification system. This has significant potential to direct clinical decisions. Patients can be selected for less aggressive treatments with greater assurance (including simple observation in some instances). At the same time, higher risk patients can be treated more appropriately with total thyroidectomy ± RAI. The identification of a molecular subgroup with a high risk of recurrence independent of tumor stage (Type 3 PTCs) is particularly impactful in this regard.

Our gene signature technology can potentially be applied to fine needle aspirates (FNA) from thyroid nodules. Currently, commercially available genomic assays for thyroid nodule aspirates (such as ThyroSeq^43^ and Thyrospec^44^) focus on diagnosis and estimating the risk of malignancy, particularly in indeterminate cell samples (Bethesda 4). Another such test, Afirma^45^, described as a genomic sequencing classifier, also detects gene variants (including BRAF^V600E^ mutations) and fusions in nodules that are clearly malignant (Bethesda 6) or are suspicious for malignancy (Bethesda 5). Our data demonstrate the limited utility of BRAF^V600E^ mutations and other genomic features for prognostication. Moreover, the transcriptomic classification we have described is biologically more informative. Being able to estimate prognosis pre-operatively would facilitate selection of patients for active surveillance, hemithyroidectomy or total thyroidectomy ± lymphadenectomy.

The three molecular subgroups that we have discovered could also potentially inform non-surgical treatment. Ras-like Type 1 PTCs are predicted to derive the greatest benefit from RAI. If future studies confirm that some Type 2 and 3 PTCs are RAI resistant, then strategies to restore the tumor’s capacity for RAI uptake are available. Currently, there are only two approved targeted therapies for patients with RAI-refractory PTC; the tyrosine kinase inhibitors sorafenib and lenvatinib. These inhibitors exert their effect by blocking the MAPK pathway ^46^. However, they have considerable toxicity and a short-lived efficacy. The biological characteristics of our molecular subtypes have uncovered future avenues for clinical research, including the use of EZH2 inhibitors and immunotherapy as primary or adjuvant therapies.

Our study is limited by a lack of prospective validation, but the clinical implications of our work to date is clear. We describe novel molecular subgroups of PTC and identify new potential therapeutic targets. We identify with specificity patient subgroups at low-risk and high-risk of disease recurrence. Most importantly, we offer a means to refine and improve clinical care in the future by outlining a diagnostic test that simplifies clinical decision making.

## AUTHOR CONTRIBUTIONS

SC, CS, FF, AH and OFB conceived and planned the experiments and clinical data analysis. CS, FF, DS, NA, AS performed the genomic and bioinformatic analyses. FF and KK created the machine learning algorithm in collaboration with OFB. YJP, CS and NA analyzed the Korean cohort. CS, MK and FK analyzed the validation cohort. CS and FK validated the targeted RNASeq assay. SC, AH, YJP and OFB reviewed the data in its clinical context. SC, CS and OFB wrote the manuscript in consultation with AH. All authors reviewed and approved the final draft of the manuscript.

## ACKNOWLEDGEMENTS

The authors wish to thank Gaurav Tripathi for his technical assistance in running the targeted RNASeq, and Manuel Machon for his analysis of data related to TDS and BRAF mutations.

## Supplementary Figure Legends

**Figure S1.** Mutation analysis summary for the three molecular subtypes. **A** Oncoplot for Type 1 tumors. **B** Oncoplot for Type 2 tumors. **C** Oncoplot for Type 3 tumors. **D.** Co-oncoplot showing the genes with significantly different mutation rates between Type 2 tumors and Type 1 tumors. **E** Co-oncoplot showing the genes with significantly different mutation rates between Type 3 tumors and Type 1 tumors (*P* < 0.05).

**Figure S2.** Summary of pathway analysis results generated using Ingenuity Pathway Analysis (IPA) of differentially expressed mRNA between molecular subtypes. **A** Canonical pathways associated with differentially expressed genes (FDR <0.05, logFC >|2|) in Type 3 tumors versus Type 1 tumors. **B** Canonical pathways associated with differentially expressed genes (FDR <0.05, logFC >|2|) in Type 2 tumors versus Type 1 tumors. P-value was calculated by IPA using the Fisher’s Exact Test.

**Figure S3.** Summary of methylation analysis of molecular subtypes. **A** Starburst plots for comparison between differential DNA methylation (x axis) and differential gene expression (y axis) when comparing Type 3 tumors with Type 1 tumors. **B** Starburst plots for comparison between differential DNA methylation (x axis) and differential gene expression (y axis) when comparing Type 2 tumors with Type 1 tumors. For **A** and **B**, the dashed black lines represent the cutoff of FDR adjusted p-values <0.05. **C.** Summary of pathway analysis results generated using Ingenuity Pathway Analysis (IPA) of differentially expressed mRNA targets of methylated genes identified by differential methylation analysis between Type 3 versus Type 1 tumors and **D** between Type 2 versus Type 1 tumors.

**Figure S4.** Summary of differential miRNA analysis between Type 3 versus Type 1 tumors. Pathway analysis results generated using Ingenuity Pathway Analysis (IPA) of differentially expressed mRNA targets of miRNAs identified by differential miRNA analysis. Negative and positive z-scores (x axis) represent predicted decreased and increased activation of a canonical pathway, respectively. The -log p-value generated by IPA represents results of a Fisher’s Exact Test analysis where p<0.05 indicates a statistically significant association.

**Figure S5.** Summary of differential miRNA analysis between Type 2 versus Type 1 tumors. Pathway analysis results generated using Ingenuity Pathway Analysis (IPA) of differentially expressed mRNA targets of miRNAs identified by differential miRNA analysis. Negative and positive z-scores (x axis) represent predicted decreased and increased activation of a canonical pathway, respectively. The -log p-value generated by IPA represents results of a Fisher’s Exact Test analysis where p<0.05 indicates a statistically significant association.

**Figure S6.** Analysis of radioactive iodine in relation to BRAF mutation, molecular subtypes, and survival. **A** Swarm plot for thyroid differentiation score in tumors with (BRAF^V600E^+) and without (BRAFV^600E^-) BRAF mutation within each molecular subtype. **B** Survival analysis revealed a significant difference in progression free survival (Log Rank p-value =0.047) between patients with the upper tertile thyroid differentiation score and the lower tertile thyroid differentiation score.

**Table S1.**
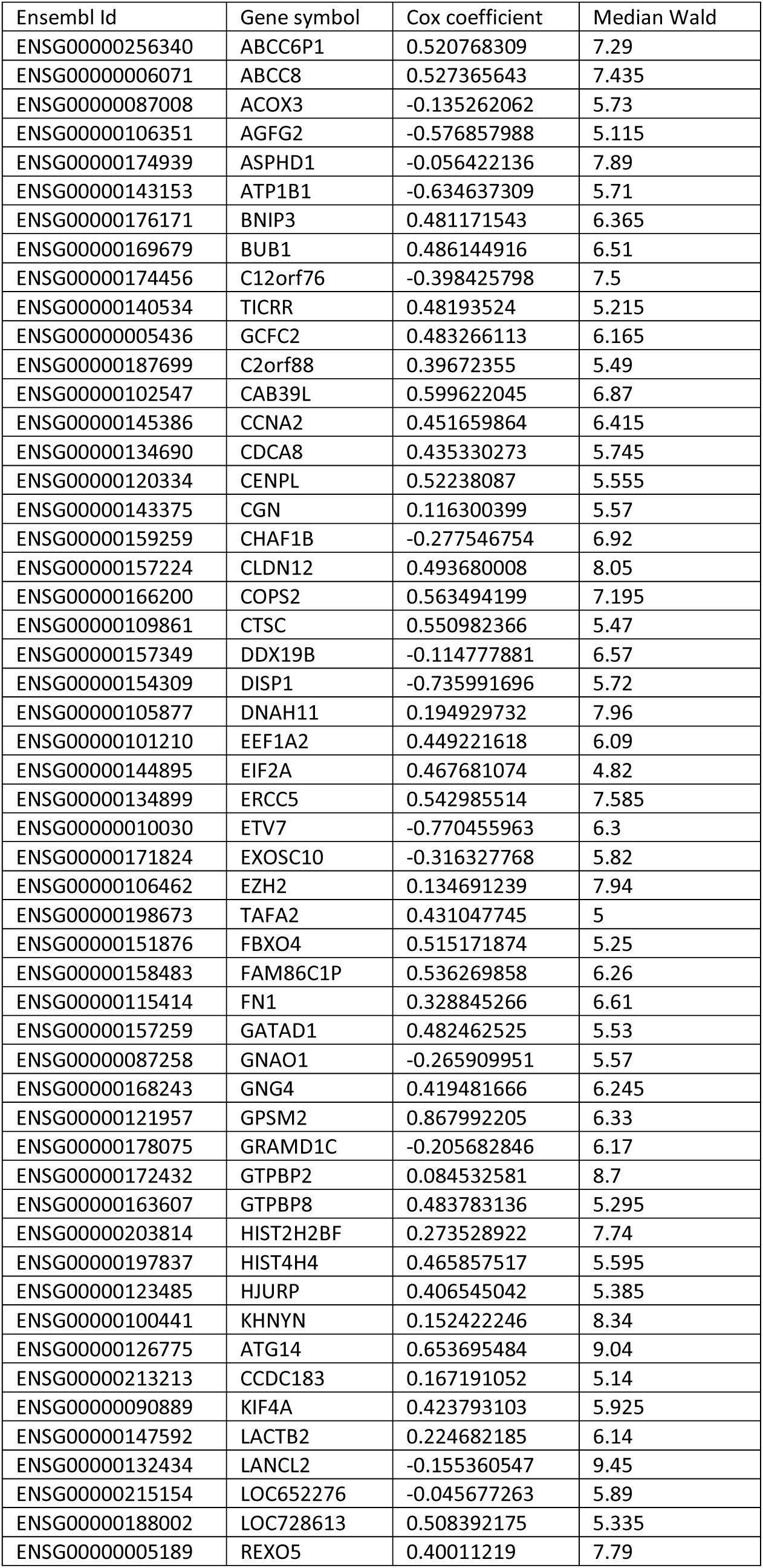
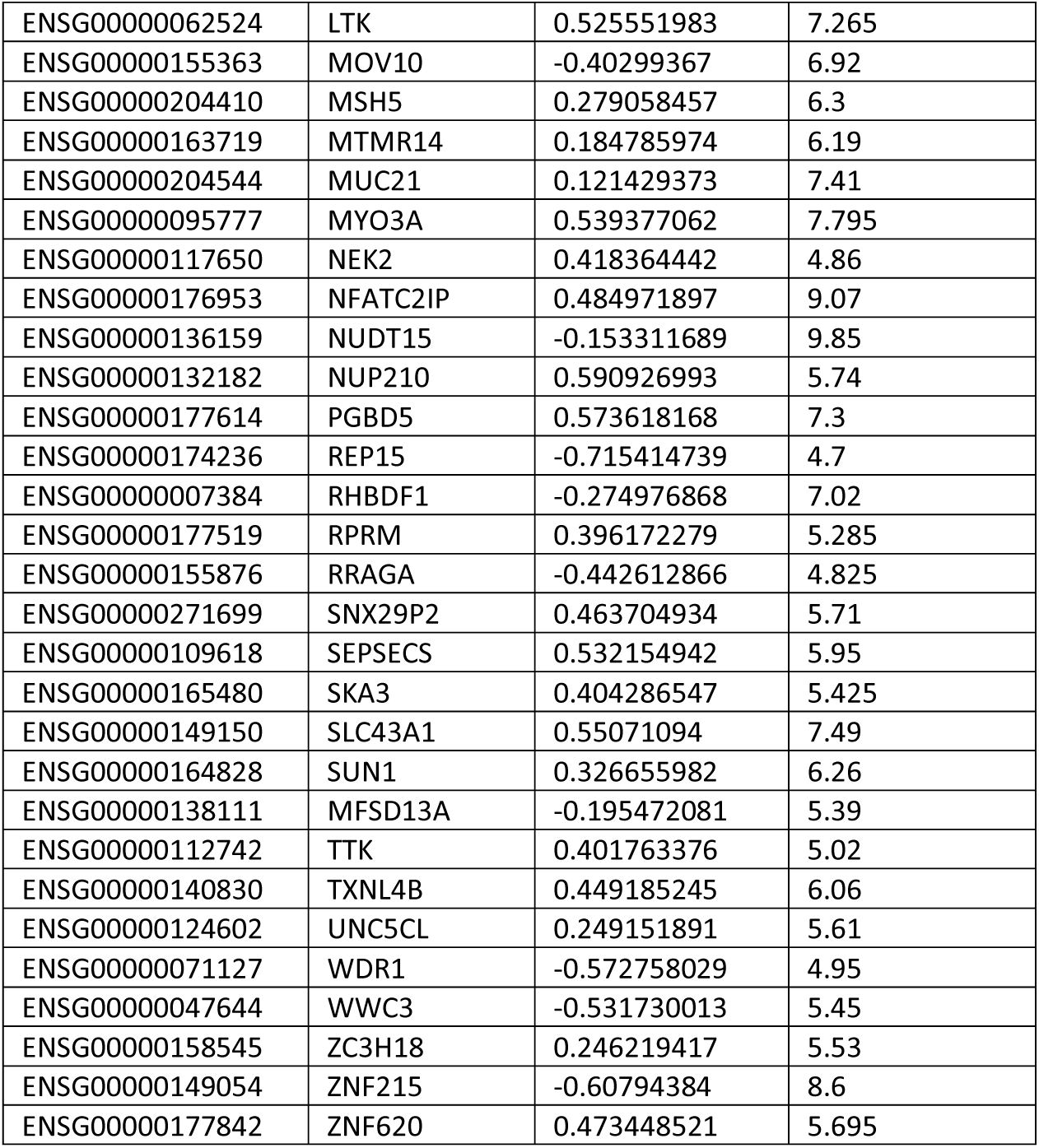
List of 82 prognostic genes identified by the HighLifeR algorithm.

**Table S2.** Univariate and multivariable analysis of factors associated with 5-year PFS in the test set.

| Risk Factor | Univariable Analysis |  | Multivariable Analysis |  |
| --- | --- | --- | --- | --- |
|  | Hazard Ratio (95 % CI) | *P* | Hazard Ratio (95 % CI) | *P* |
| Sex (female v. male) | 1.37 (0.63 – 2.94) | 0.43 |  |  |
| Age (< 50 years v. ≥50 years) | 1.37 (0.66 – 2.86) | 0.40 |  |  |
| Tumor size (≤4 cm v. >4 cm) | 1.39 (0.64 – 3.01) | 0.40 |  |  |
| Histological type |  |  |  |  |
| Classical | 1 (reference) |  |  |  |
| Follicular | 0.47 (0.14 – 1.58) | 0.23 |  |  |
| Tall Cell | 0.92 (0.28 – 3.05) | 0.89 |  |  |
| Other | 0.00 (0.00 – inf) | 1.0 |  |  |
| T stage |  |  |  |  |
| T1 | 1 (reference) |  | 1 (reference) |  |
| T2 | 3.15 (0.64 – 15.61) | 0.16 | 4.33 (0.86 – 21.86) | 0.08 |
| T3 | 5.16 (1.19 – 22.34) | **0.03** | 4.49 (0.98 – 20.58) | 0.053 |
| T4 | 11.80 (2.15 – 64.71) | **0.004** | 3.27 (0.43 – 24.97) | 0.25 |
| N stage (N0/NX v. N1) | 1.85 (0.86 – 3.98) | 0.12 |  |  |
| M stage (M0 v. M1) | 162.50 (10.16 – 2598) | **3.19 x 10<sup>-4</sup>** | 135.68 (6.49 – 2836.73) | **1.55 x 10<sup>-3</sup>** |
| BRAF<sup>V600E</sup> mutation (Absent v. Present) | 1.18 (0.57 – 2.45) | 0.66 |  |  |
| TERT promoter mutation (Absent v. Present) | 2.05 (0.71 – 5.90) | 0.18 |  |  |
| Thyroid Differentiation Score |  |  |  |  |
| < -1 | 1 (reference) |  |  |  |
| -1 to 0 | 0.44 (0.15 – 1.31) | 0.14 |  |  |
| > 0 | 0.33 (0.10 – 1.08) | 0.07 |  |  |
| BRAF<sup>V600E</sup>-RAS class |  |  |  |  |
| BRAF-like | 1 (reference) |  |  |  |
| RAS-like | 0.56 (0.16 – 1.93) | 0.36 |  |  |
| ATA Risk |  |  |  |  |
| Low | 1 (reference) |  | 1 (reference) |  |
| Intermediate | 2.19 (0.91 – 5.26) | 0.08 | 1.31 (0.47 – 3.60) | 0.60 |
| High | 3.82 (1.55 – 9.41) | **3.62 x 10<sup>-3</sup>** | 3.12 (0.91 – 10.79) | 0.07 |
| Molecular Subtype |  |  |  |  |
| Type 2 | 1 (reference) |  | 1 (reference) |  |
| Type 1 | 0.69 (0.19 – 2.52) | 0.58 | 0.93 (0.24 – 3.62) | 0.92 |
| Type 3 | 2.75 (1.25 – 6.08) | **0.01** | 3.64 (1.55 – 8.56) | **3.06 x 10<sup>-3</sup>** |

HR = Hazard ratio, the ratio of recurrence rates in two comparator groups for any independent variable (for binary variables) or the logarithm of the change in death rate per unit change of the independent variable (if the variable is continuous).
CI = Confidence interval

**Table S3.**
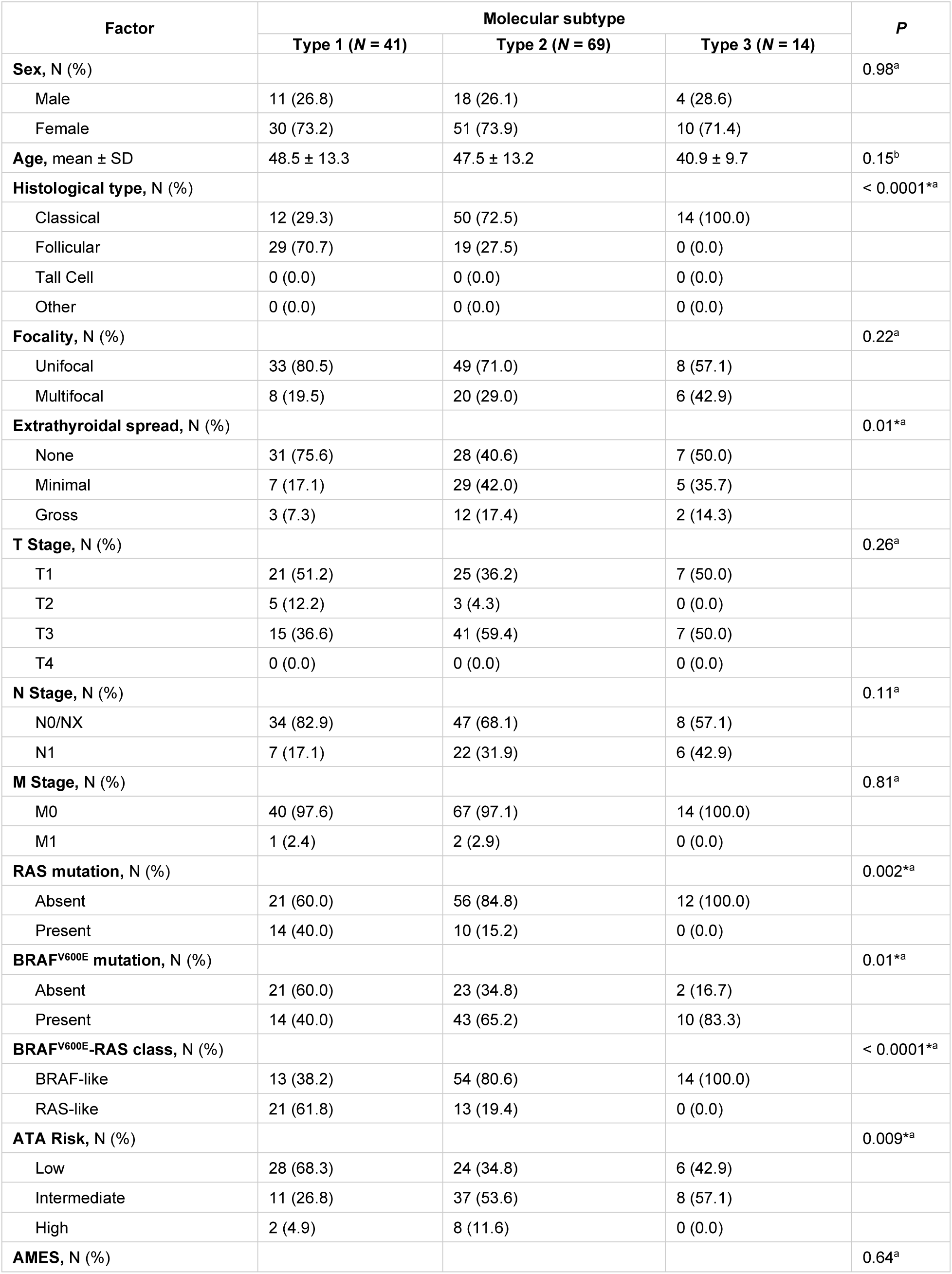
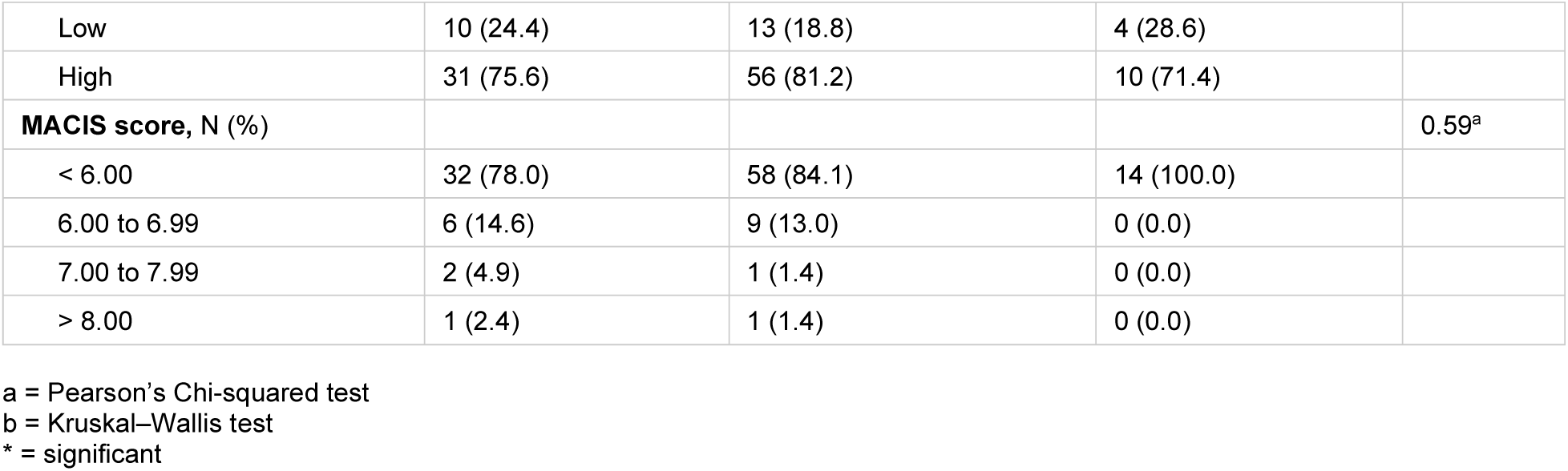
Clinical characteristics of molecular subtypes assigned to the external validation dataset from Korea.

| Factor | Molecular subtype |  |  | P |
| --- | --- | --- | --- | --- |
|  | Type 1 (N = 41) | Type 2 (N = 69) | Type 3 (N = 14) |  |
| **Sex, N (%)** |  |  |  | 0.98<sup>a</sup> |
| Male | 11 (26.8) | 18 (26.1) | 4 (28.6) |  |
| Female | 30 (73.2) | 51 (73.9) | 10 (71.4) |  |
| **Age, mean ± SD** | 48.5 ± 13.3 | 47.5 ± 13.2 | 40.9 ± 9.7 | 0.15<sup>b</sup> |
| **Histological type, N (%)** |  |  |  | < 0.0001<sup>*a</sup> |
| Classical | 12 (29.3) | 50 (72.5) | 14 (100.0) |  |
| Follicular | 29 (70.7) | 19 (27.5) | 0 (0.0) |  |
| Tall Cell | 0 (0.0) | 0 (0.0) | 0 (0.0) |  |
| Other | 0 (0.0) | 0 (0.0) | 0 (0.0) |  |
| **Focality, N (%)** |  |  |  | 0.22<sup>a</sup> |
| Unifocal | 33 (80.5) | 49 (71.0) | 8 (57.1) |  |
| Multifocal | 8 (19.5) | 20 (29.0) | 6 (42.9) |  |
| **Extrathyroidal spread, N (%)** |  |  |  | 0.01<sup>*a</sup> |
| None | 31 (75.6) | 28 (40.6) | 7 (50.0) |  |
| Minimal | 7 (17.1) | 29 (42.0) | 5 (35.7) |  |
| Gross | 3 (7.3) | 12 (17.4) | 2 (14.3) |  |
| **T Stage, N (%)** |  |  |  | 0.26<sup>a</sup> |
| T1 | 21 (51.2) | 25 (36.2) | 7 (50.0) |  |
| T2 | 5 (12.2) | 3 (4.3) | 0 (0.0) |  |
| T3 | 15 (36.6) | 41 (59.4) | 7 (50.0) |  |
| T4 | 0 (0.0) | 0 (0.0) | 0 (0.0) |  |
| **N Stage, N (%)** |  |  |  | 0.11<sup>a</sup> |
| N0/NX | 34 (82.9) | 47 (68.1) | 8 (57.1) |  |
| N1 | 7 (17.1) | 22 (31.9) | 6 (42.9) |  |
| **M Stage, N (%)** |  |  |  | 0.81<sup>a</sup> |
| M0 | 40 (97.6) | 67 (97.1) | 14 (100.0) |  |
| M1 | 1 (2.4) | 2 (2.9) | 0 (0.0) |  |
| **RAS mutation, N (%)** |  |  |  | 0.002<sup>*a</sup> |
| Absent | 21 (60.0) | 56 (84.8) | 12 (100.0) |  |
| Present | 14 (40.0) | 10 (15.2) | 0 (0.0) |  |
| **BRAF<sup>V600E</sup> mutation, N (%)** |  |  |  | 0.01<sup>*a</sup> |
| Absent | 21 (60.0) | 23 (34.8) | 2 (16.7) |  |
| Present | 14 (40.0) | 43 (65.2) | 10 (83.3) |  |
| **BRAF<sup>V600E</sup>-RAS class, N (%)** |  |  |  | < 0.0001<sup>*a</sup> |
| BRAF-like | 13 (38.2) | 54 (80.6) | 14 (100.0) |  |
| RAS-like | 21 (61.8) | 13 (19.4) | 0 (0.0) |  |
| **ATA Risk, N (%)** |  |  |  | 0.009<sup>*a</sup> |
| Low | 28 (68.3) | 24 (34.8) | 6 (42.9) |  |
| Intermediate | 11 (26.8) | 37 (53.6) | 8 (57.1) |  |
| High | 2 (4.9) | 8 (11.6) | 0 (0.0) |  |
| **AMES, N (%)** |  |  |  | 0.64<sup>a</sup> |
| Low | 10 (24.4) | 13 (18.8) | 4 (28.6) |  |
| High | 31 (75.6) | 56 (81.2) | 10 (71.4) |  |
| **MACIS score, N (%)** |  |  |  | **0.59<sup>a</sup>** |
| < 6.00 | 32 (78.0) | 58 (84.1) | 14 (100.0) |  |
| 6.00 to 6.99 | 6 (14.6) | 9 (13.0) | 0 (0.0) |  |
| 7.00 to 7.99 | 2 (4.9) | 1 (1.4) | 0 (0.0) |  |
| > 8.00 | 1 (2.4) | 1 (1.4) | 0 (0.0) |  |

a = Pearson's Chi-squared test
b = Kruskal–Wallis test
\* = significant

**Table S4.** Five-year recurrence rates for different cohorts.

| Cohort | Molecular subtype |  |  |
| --- | --- | --- | --- |
|  | Type 1 | Type 2 | Type 3 |
| **TCGA Discovery** |  |  |  |
| Early | 3/54 (5.6%) | 1/73 (1.4%) | 5/34 (14.7%) |
| Advanced | 5/28 (17.9%) | 1/82 (1.2%) | 7/52 (13.5%) |
| All | 8/82 (9.8%) | 2/155 (1.3%) | 12/86 (14.0%) |
| **TCGA Test** |  |  |  |
| Early | 1/22 (4.5%) | 1/28 (3.6%) | 4/12 (33.3%) |
| Advanced | 2/15 (13.3%) | 8/49 (16.3%) | 12/37 (32.4%) |
| All | 3/37 (8.1%) | 9/77 (11.7%) | 16/49 (32.7%) |
| **Korea cohort** |  |  |  |
| Early | 0/32 (0%) | 0/46 (0%) | 0/8 (0%) |
| Advanced | 0/9 (0%) | 1/23 (4.3%) | 0/6 (0%) |
| All | 0/41 (0%) | 1/69 (1.4%) | 0/14 (0%) |
| **Edmonton cohort** |  |  |  |
| Early | 0/11 (0%) | 0/18 (0%) | 2/12 (16.7%) |
| Advanced | 0/11 (0%) | 13/51 (25.5%) | 5/29 (17.2%) |
| All | 0/22 (0%) | 13/69 (18.8%) | 7/41 (17.1%) |
| **All cohorts combined** |  |  |  |
| Early | 4/119 (3.4%) | 2/165 (1.2%) | 11/66 (16.7%) |
| Advanced | 7/63 (11.1%) | 23/205 (11.2%) | 24/124 (19.4%) |
| All | 11/182 (6.0%) | 25/370 (6.8%) | 35/190 (18.4%) |

## METHODS

### Patients and transcriptional data for prognostic model

To train and validate the prognostic classifier, we used the RNA-seq dataset from PTC primary tumor samples. These are available in the Cancer Genome Atlas (TCGA) consortium database (https://portal.gdc.cancer.gov/). Details of the patient inclusion criteria, sites of sample collection, and transcriptional analysis methodology can be found elsewhere ^9^. The data were randomly split into thirds. Two thirds (*N* = 335) were assigned to be the discovery dataset and the remaining third was assigned to be the test set (*N* =167). Additional accompanying TCGA data (e.g., mutation data and copy number variation data) were downloaded using the TCGAbiolinks package (version 2.19.0)^47^ and from Genomic Data Commons^48^. Harmonized molecular data were used (aligned to hg38).

Two-tailed Student’s *t-*tests, ANOVA tests, correlations, Fisher’s Exact tests, Pearson Chi-squared tests and McNemar’s test were conducted using IBM-SPSS v.28.0 statistical software (IBM). *P* and n values are indicated in the figure legends or in the figures themselves. The R package smoothROCtime was used for time-dependent ROC curve estimations^49^. Unless otherwise stated, *P* < 0.05 was considered significant.

### Identification of prognostic genes

*HighLifeR* was designed to address the intrinsic problems with Cox Proportional Hazard analysis in highly dimensional genomic datasets. Specifically, the Cox method requires a minimum number of events per variable studied, and has limited capacity filtering of the most impactful, yet non-parallel, variables (genes), limiting its suitability for multivariable analysis of high-dimension genomic datasets. *HighLifeR* leverages the Partial Cox Regression method of Li and Gui ^50^. In the *HighLifeR* algorithm, the multivariable Cox proportional hazards are estimated through recursive generation of predictive latent components. This was done using a partial least squares (PLS) extension on survival data by testing millions of combinations of genes and patients in a regulated machine learning environment. The training population is also randomized into “virtual cohorts” over the course of 20 rounds of testing, each composed of at least 70% of patients, with resampling. This approach limits the effect from outliers and substantially reduces the dataset dimensionality, and it can be used to generate a PC-R based model for predicting survival outcomes. It is applicable when prior knowledge is limited, such as in genomic studies. At the same time, it is possible to adjust for known prognostic factors by stratifying the virtual cohorts.

Three main components of the *HighLifeR* statistical pipeline include: 1. massively-parallel combinatorial rounds to establish the prognostic associations for each gene in the variable space; 2. selection of variables (genes) with the highest potential from step 1 for the purpose of developing an accurate and comprehensive, yet parsimonious, prognostic classifiers; and 3. construction of a composite prognostic scoring model. Wide implementation of randomization procedures in the pipeline, including sample allocation to test and validation sets and in combination rounds, reduces the possibility of detecting a sample set-dependent prognostic profile. To select prognostic genes, we required genes to be consistently ranked among the top 200 genes based on their association with time to recurrence and have a prognostic impact (Wald statistic) greater than the half of the maximum Wald statistic in our training set (>4.7).

### Identification of risk classes

We identified prognostic genes using HighLifeR and then conducted unsupervised clustering in the discovery dataset to see how patients were grouped based on the expression of these genes. We then created Kaplan-Meier survival curves to compare groups identified by unsupervised clustering and tested for group differences (Log Rank Test) using IBM-SPSS v.28.0 statistical software (IBM). Once clusters in the training set were determined, genes in the clusters were used to build the prognostic classifiers. These gene clusters were then compared using the test dataset. Prognostic model development and testing were conducted using the WEKA software suite (version 3.8.4)^51^. Heatmaps and unsupervised clustering were conducted in R using the ComplexHeatmap package (version 2.10.0)^52^.

### Differential mutation

Raw maf files containing mutation data were analyzed. Oncoplots and coOncoplots were created for visualization using the R package maftools (version 2.6.05)^53^. Differential mutation was also conducted using maftools which performs Fisher’s test to compare mutations frequency between two groups.

### Differential expression

RNA-seq data (HT-Seq counts) were downloaded, prepared, and normalized using TCGAbiolinks. Differential expression was conducted using the EdgeR method^54^. The low-risk group served as the reference group. We used log2 Fold Change >1 and adjusted p values <0.05 to identify differentially expressed genes. mRNA functional analysis included submitting differentially expressed genes to IPA (QIAGEN Inc., https://www.qiagenbioinformatics.com/products/ingenuitypathway-analysis) and Gene Set Enrichment (GSEA)^55^ analysis using the 50 molecular signatures called the Hallmark gene sets.

### Differential methylation

Beta values (Infinium Human Methylation 450k platform) were downloaded using the GDC Data Transfer Tool User (version 1.6.1). Somatic differentially methylated CpGs analysis was conducted using the TCGAanalyze_DMC function in TCGAbiolinks. Significance cut-offs were an adjusted p-value of <0.05 and a mean difference in beta values > 0.2. Once differentially DNA methylated genes were identified, integration of the differentially expressed genes allowed us to explore genes which were hyper-methylated and down-regulated and/or hypo-methylated and up-regulated in each risk group compared with the low-risk group (which always served as the reference group). A starburst plot was created for each risk comparison for visualization of the methylation and gene expression relationships. In the starburst plots, the p-values are multiplied by the sign of difference of beta values. The dashed black lines indicate the p-value at 0.05. The function of the genes highlighted in the starburst plots was examined manually or through using IPA software.

### Differential miRNA

BCGSC miRNA Profiling Pipeline data were downloaded using TCGAbiolinks. Differentially expressed miRNA were identified using the DESeq2 package^56^. miRNA-gene pairs were identified which corresponded to functional validation publications reported by MiRTarBase (version 9.0)^57^, for stronger (luciferase reporter, qPCR, western blot) and weaker experimental evidence types. Differentially downregulated genes were paired to differentially upregulated miRNA, and differentially upregulated genes were paired to differentially downregulated miRNA. These pairs were submitted to IPA for pathway analysis.

### Immune cell infiltration analysis

Immune cell fractions were estimated using the mixed sample gene expression deconvolution algorithm CIBERSORT^58^. Using a set of 22 immune cell reference expression profiles (LM22 signature matrix), the relative abundance of immune cells in the tissue were estimated. The overall fraction of each immune cell type in the tissue was calculated by multiplying the CIBERSORT relative abundance in the leukocyte fraction (LF), as explained elsewhere^59^.

The T cell-inflamed gene expression profile (GEP) was derived across a variety of solid tumors. It is composed of 18 inflammatory genes related to antigen presentation, chemokine expression, cytolytic activity, and adaptive immune resistance, including CCL5, CD27, CD274 (PD-L1), CD276 (B7-H3), CD8A, CMKLR1, CXCL9, CXCR6, HLA-DQA1, HLA-DRB1, HLA-E, IDO1, LAG3, NKG7, PDCD1LG2 (PDL2), PSMB10, STAT1, and TIGTT ^60^. The GEP scores were calculated as a weighted sum of normalized expression values for the 18 genes listed above.

### External validation set

To validate our prognostic model, we accessed another public dataset containing RNA-Seq data from PTC tumors from 124 Korean patients with PTC (PRJEB11591)^22^. Details regarding sequencing can be found elsewhere^22^. FASTQ files were accessed from the European Nucleotide Archive using BaseSpace (https://basespace.illumina.com/home/indexIllumina, San Diego CA) and were subsequently analyzed using FASTQC for quality control (https://www.bioinformatics.babraham.ac.uk/projects/fastqc). Salmon was used to quantify transcript abundance from RNA-Seq reads on the Galaxy web interface^61, 62^. TPM counts were scaled prior to prognostication using our prediction model.

### Targeted RNASeq for Molecular Classification of Clinical Samples

This analysis was approved the Health Research Ethics Board of Alberta Cancer Committee (Ethics no. HREBA.CC-18-0285). Fresh frozen papillary thyroid carcinoma samples (*N* = 136) were acquired from the tumor bank in Edmonton, Canada and stored at −80C prior to analysis. RNA was extracted using the RNeasy Mini Kit (Qiagen) according to manufacturer’s instructions. The integrity of RNA was determined by electrophoresis using 2100 Bioanalyzer (Agilent Technologies).

A custom panel was designed using probes for the 82 prognostic genes and 10 internal controls. Internal controls were selected based on their low expression variance between tumors; 5 were high abundance genes, and 5 were low abundance genes. External RNA controls developed by the External RNA Controls Consortium (ERCC; Invitrogen/Thermo Fisher) composed of 92 unique transcripts derived and traceable from NIST-certified DNA plasmids were added for technical quality control.

cDNA synthesis and library preparation were performed according to the Illumina RNA prep with enrichment (L) tagmentation protocol. The prepared libraries were pooled and sequenced on the Illumina MiSeq platform using the custom panel designed using the HighLifeR prognostic genes, the internal controls and the ERCC transcripts. MiSeq Reagents Kit v3 (Illumina) were used according to the manufacturer’s instructions. Sequenced reads were quality control (QC) checked using FastQC (version 0.11.9), trimmed using Fastp (version 0.23.2), and, quantified using Salmon (1.5.1)^61^ using the quasi-mapping mode. FastQC, Fastp and Salmon were run on Galaxy (version 22.05) Transcript level counts were summarized to gene level using tximport (version 1.24.0). Gene level data were used for the prediction using the classification models designed to classify tumors as Type 1, Type 2, or Type 3.

