## Supplemental Figures 1-6 for "A Clinically Useful and Biologically Informative Genomic Classifier for Papillary Thyroid Cancer"

**Supplementary Figure S1**

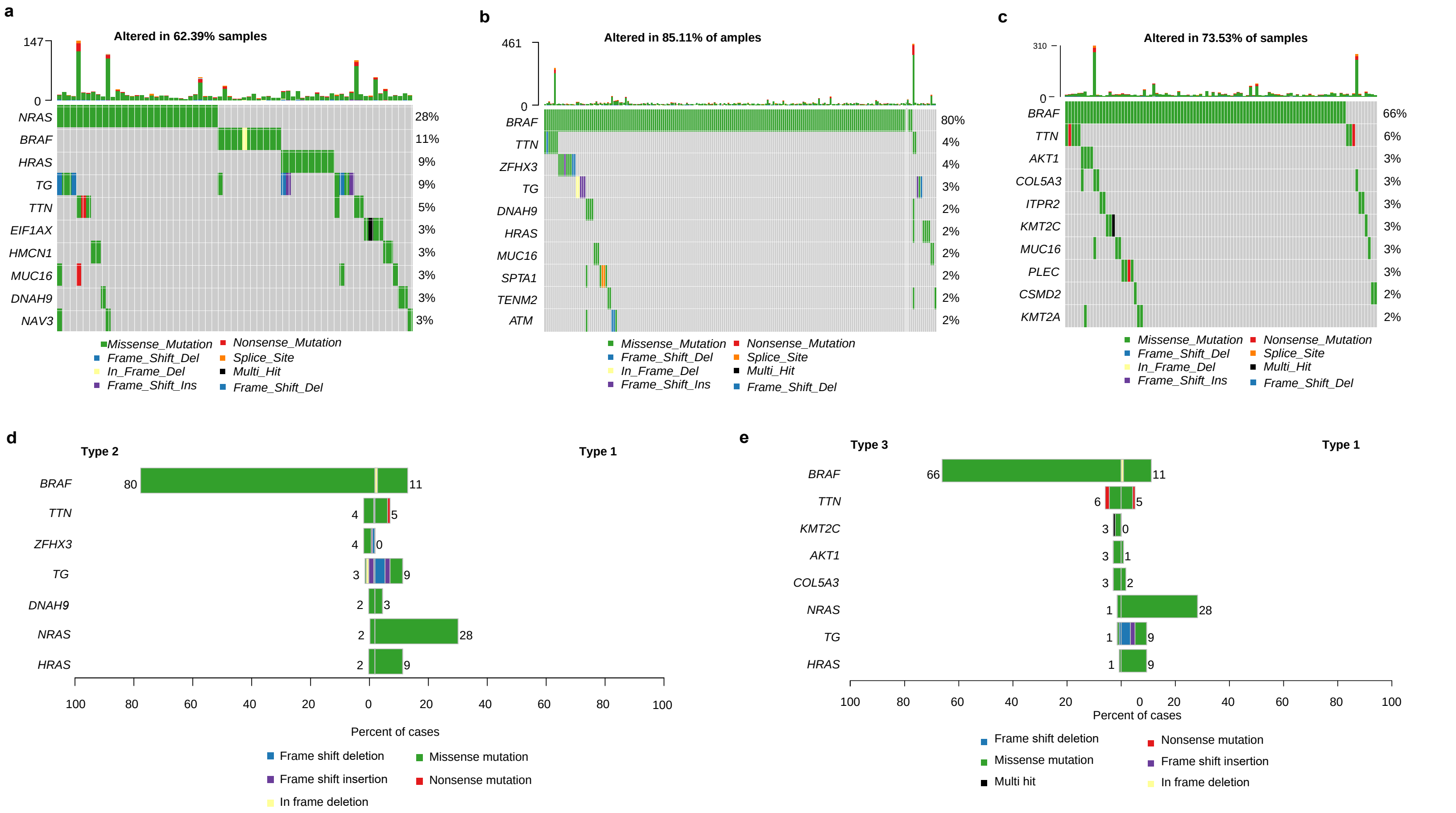

Supplementary Figure S2

a

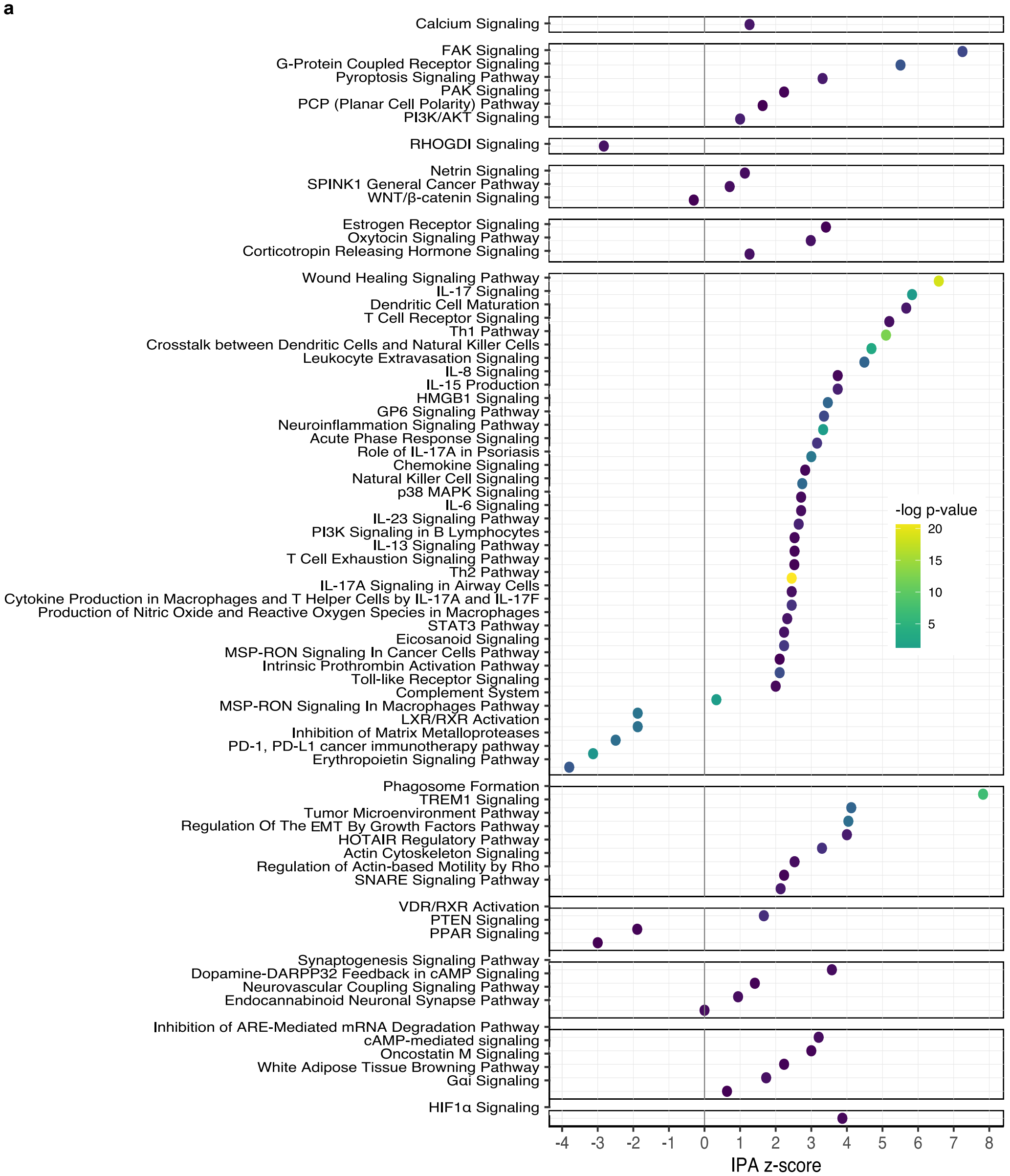


b

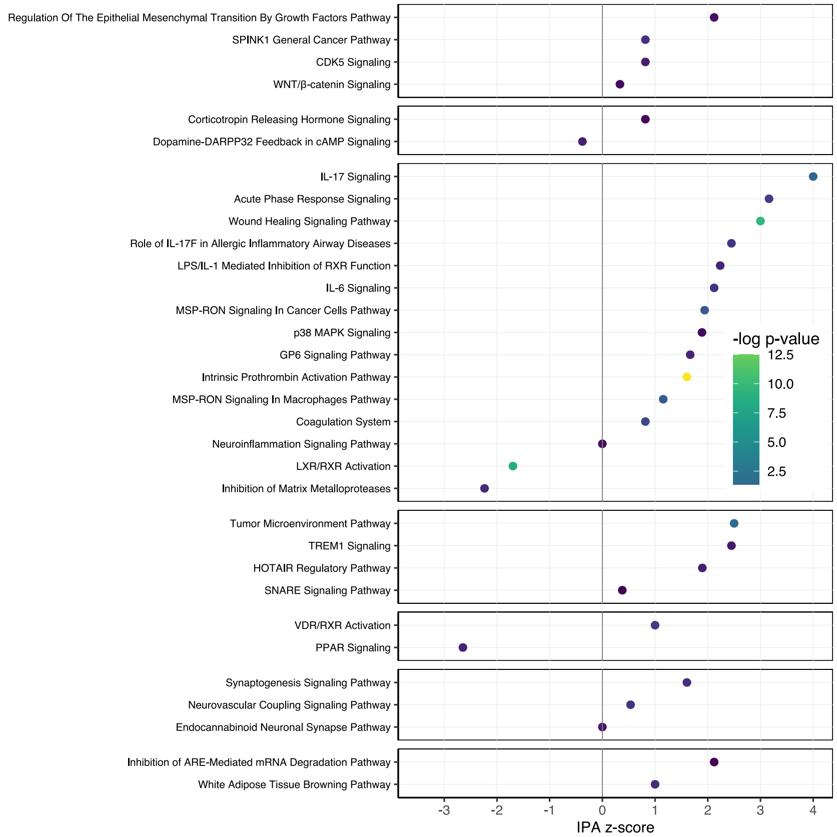

**Supplementary Figure S3**

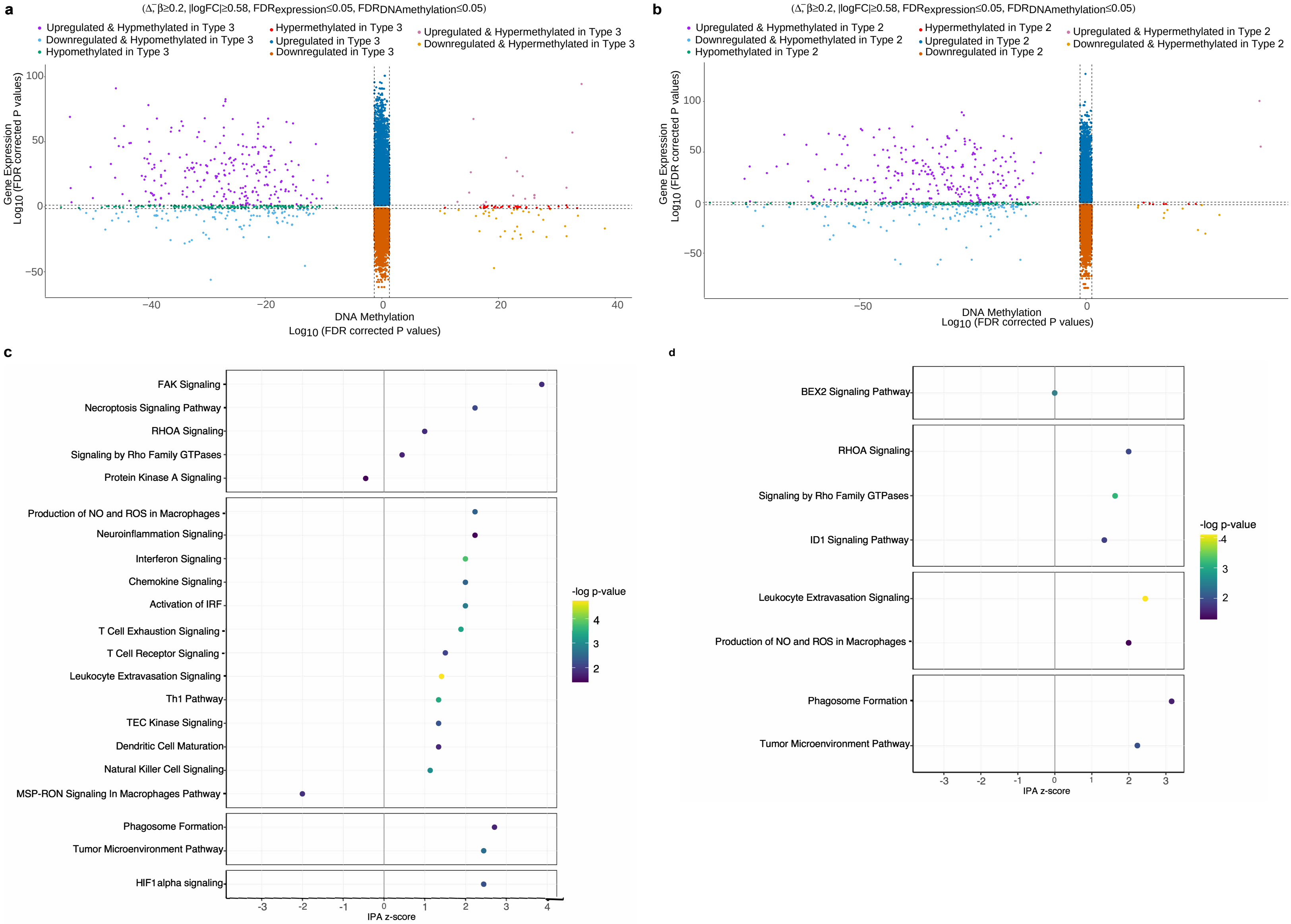

**Supplementary Figure S4**

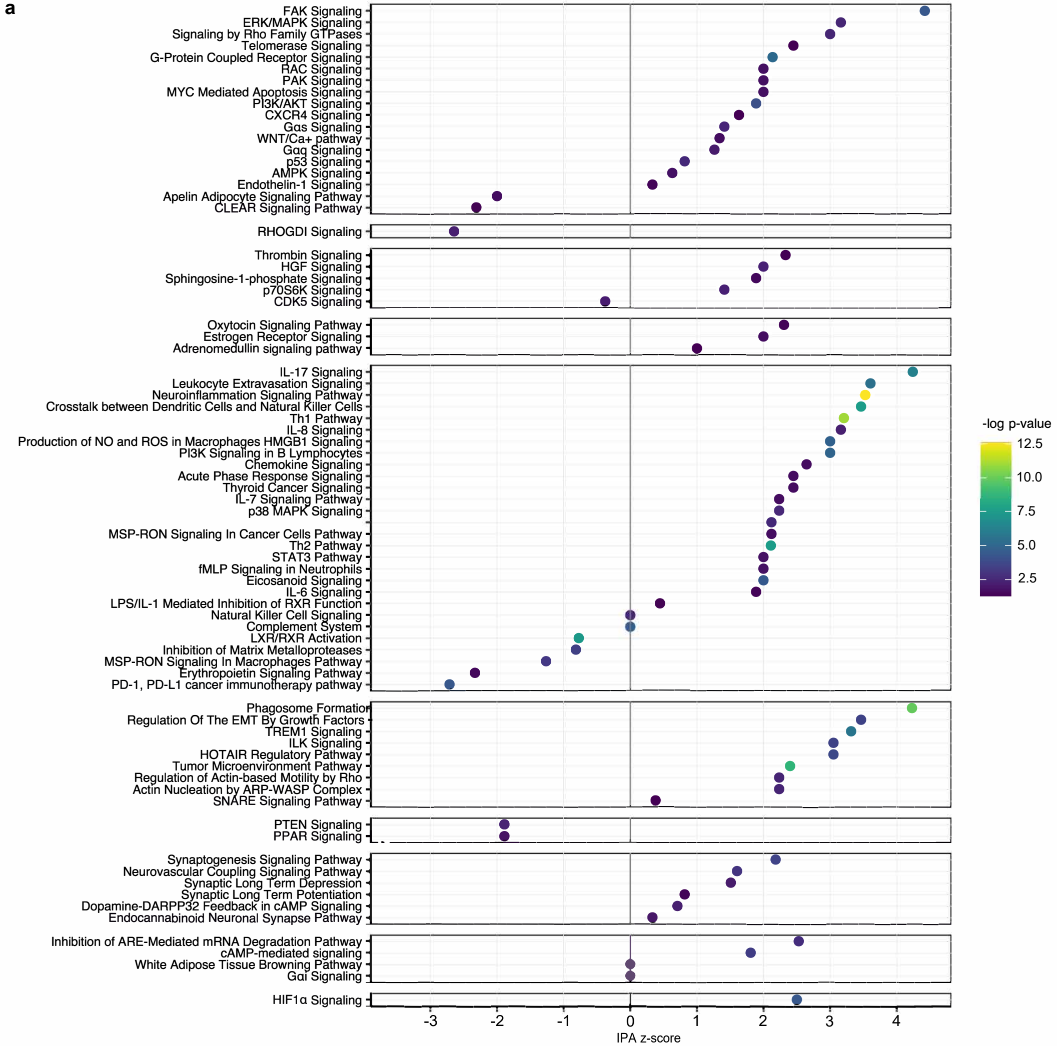

Supplementary Figure S5

a

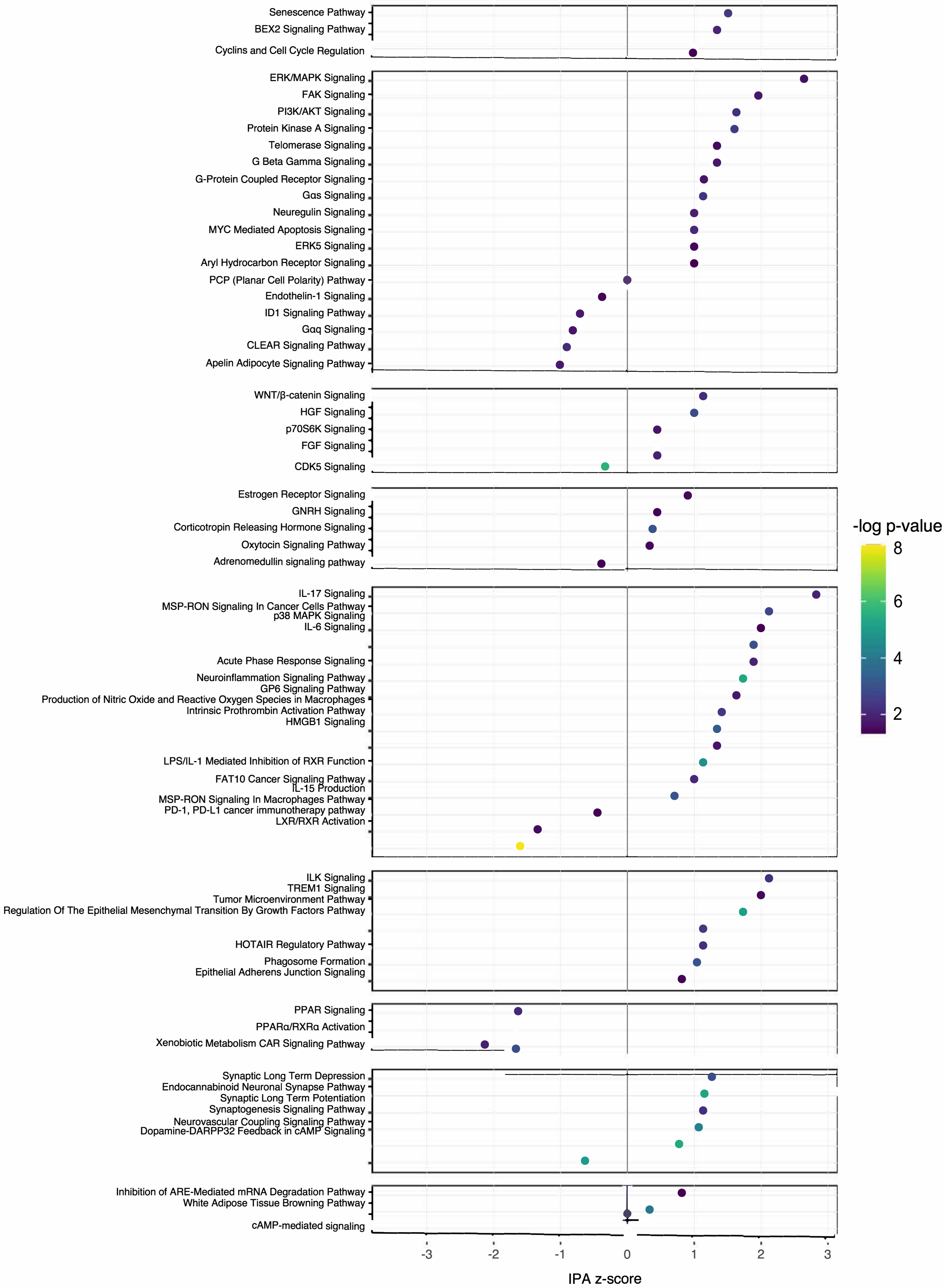

**Supplementary Figure S6**

a

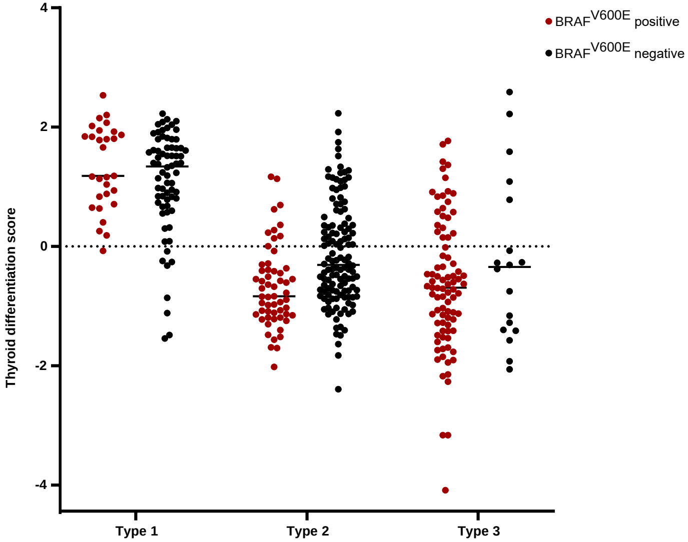


b

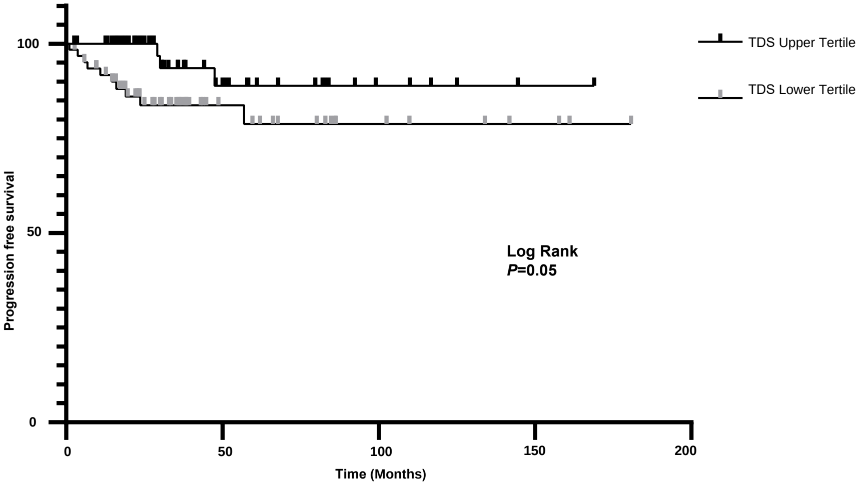
